# CryoEM structure of QacA, an antibacterial efflux transporter from *Staphylococcus aureus*

**DOI:** 10.1101/2022.07.09.499445

**Authors:** Puja Majumder, Shahbaz Ahmed, Pragya Ahuja, Arunabh Athreya, Rakesh Ranjan, Aravind Penmatsa

## Abstract

Efflux of antibacterial compounds is a major mechanism for developing antimicrobial resistance. In the Gram-positive pathogen *Staphylococcus aureus*, QacA, a 14 transmembrane (TM) helix containing major facilitator superfamily antiporter, mediates proton-coupled efflux of mono and divalent cationic antibacterial compounds. In this study, we report the cryoEM structure of QacA, with a single mutation D411N that improves homogeneity and retains efflux activity against divalent cationic compounds like dequalinium and chlorhexidine. The structure of substrate-free QacA, complexed to two single-domain camelid antibodies, was elucidated to a resolution of 3.6 Å. The structure displays an outward-open conformation with an extracellular hairpin loop, which is conserved in a subset of DHA2 transporters and its deletion causes a loss of function in the transporter. Modeling and simulations of QacA’s cytosol-facing and occluded conformations reveal asymmetry in the rocker-switch mode of QacA’s conformational shifts, providing new insights into the organization and structural dynamics of DHA2 members.

## Introduction

Pathogenic bacteria gain antimicrobial resistance (AMR) through diverse mechanisms to counter the biocidal effects of antibiotics and antibacterial compounds. One of the major routes of developing AMR is through the process of active efflux of biocides toxic to the pathogen^1^. A diverse array of pumps and transporters are employed by drug-resistant bacteria to efflux antibiotics or antibacterial compounds^2,3^. Besides being involved in direct efflux, efflux pumps and transporters are known to enhance the persistence among bacterial populations carrying their genes^4^. A wide range of drug-resistant pathogens express diverse efflux pump machinery within their membranes that includes the MFS (major facilitator superfamily), MATE (Multi-drug and toxin extrusion family), SMR (small multidrug resistance), ABC (ATP-binding cassette), RND (Resistance-Nodulation Division) and PACE (proteobacterial antimicrobial compound efflux) family transporters of which the RND and PACE family transporters are primarily observed in Gram-negative pathogens^3^.

Among the numerous pathogens that display AMR, the WHO has annotated a priority list of twelve drug-resistant pathogenic bacteria that require novel antibiotics to counter them^5^. The list annotates the Gram-positive pathogen, *Staphylococcus aureus*, as a high-priority pathogen that causes a wide array of localized or systemic infections, including bacteremia, endocarditis, and implant infections^6^. Drug-resistant strains of *Staphylococcus aureus* express a diverse set of MFS antiporters in their membranes, including the chromosomally encoded transporters NorA, NorB, and NorC that provide resistance against fluoroquinolones^7^, and QacA and QacB that are plasmid-encoded and are involved in antibacterial efflux^8,9^. QacA is highly prevalent in *S*.*aureus* strains resistant to cationic antibacterial compounds, particularly among those associated with nosocomial infections^10^. The MFS transporters involved in drug:proton antiport (DHA) are divided into DHA1 and DHA2 depending on the number of transmembrane (TM) helices (12 or 14, respectively) in each transporter^11^. While transporters like NorA, MdfA, and LmrP comprise 12 TM helices and belong to the DHA1 class of transporters^12-14^, QacA, LfrA, and SmvA comprise 14 TM helices and are part of the DHA2 family^9,15,16^.

QacA is a prototypical DHA2 member and is the first proton-coupled antibacterial efflux transporter to be functionally characterized^17^. Its expression is regulated by the repressor, QacR, which works as a negative regulator and allows *qacA* transcription in response to interactions with quaternary ammonium compounds (QACs)^18,19^. QacA is a promiscuous antiporter capable of transporting nearly thirty different cationic antibacterial compounds^20^. Its efflux ability encompasses divalent and monovalent cationic compounds, including commonly used antibacterials like benzalkonium chloride, dequalinium, and chlorhexidine^20^. A homology model of QacA is observed to employ a set of six residues with acidic side chains (D34, D61, D323, E406, E407, and D411) that surround the solvent-accessible vestibule of the transporter^21^. They play diverse roles through their involvement in competitive substrate recognition or undergo protonation to facilitate enhanced stoichiometry for drug:proton antiport^21^. The experimentally determined topology of QacA also depicts the presence of a large extracellular loop between TMs 13 and 14, whose role remains unexplored. Although a structure of a minimally characterized DHA2 member, NorC, is available^22^, QacA represents a prototypical DHA2 member with its ability to transport cationic antibacterials, whereas the substrates of NorC remain to be identified^22^.

This study delves into cryoEM structure elucidation of QacA using a single mutant of D411N that enhances its homogeneity. We also employ two Indian camelid antibodies (ICabs A4 and B7) as fiducial markers to obtain the first high-resolution structure of QacA in an outward-open state at a resolution of 3.6 Å. The structure provides insights into the positions of the protonatable acidic residues around the solvent-accessible vestibule. It identifies the extracellular loop 7 as a structured hairpin loop connecting TMs 13 and 14 that could influence substrate efflux in QacA. This motif is conserved in a subset of DHA2 transporters; based on molecular dynamics (MD) simulation trajectories we predict that this motif interacts with extracellular loop 1 on the N-terminal domain, likely playing a structural role during the transport cycle. Based on simulations and models of QacA’s alternate conformations, we suggest the presence of asymmetry in the rocker-switch mechanism in related DHA2 members.

## Results

### Construct optimization to enhance QacA homogeneity

The QacA_WT_ was heterologously expressed and isolated from the membranes of an *E*.*coli* strain lacking the efflux pumps *acrB, mdfA* and *ydhE*. Despite optimizing detergent conditions and identifying undecyl–β-D-maltoside as the optimal detergent for purification, the QacA_WT_ construct displays significant levels of aggregation in size exclusion chromatography (Fig.1a). As part of the analysis of protonation sites within QacA that affect substrate efflux, we identified QacA_D411N_ as a mutant that significantly improves the sample’s homogeneity to facilitate structural studies (Fig. 1b). We observed in our earlier studies that D411 is essential for the transport of monovalent cations like ethidium (Et) and tetraphenyl phosphonium (TPP^+^), yet QacA_D411N_ retains the ability to transport some divalent cationic substrates like chlorhexidine and dequalinium, which are popular antibacterial compounds (Fig. 1c)^21^. Conformational stabilization through cytosolic rim residue mutagenesis had been successfully attempted in the case of MdfA^23^; hence we sought to further stabilize QacA_D411N_ by introducing additional mutants of polar residues at the cytosolic membrane interface, including E137Q (TM4-TM5), E141Q, R142Q and A143N (TM5), D267N (TM8-TM9), L389N and L392N (TM12) (Supplementary Fig. 1a, b). Of these, QacA_D411N/E137Q_ displayed a mildly weakened efflux, suggesting a lowered ability to cycle through conformational transitions. The construct was purified to homogeneity (Supplementary Fig. 1c) and used as the antigen for immunizing camels to raise single-domain camelid antibodies.

**Fig. 1.**
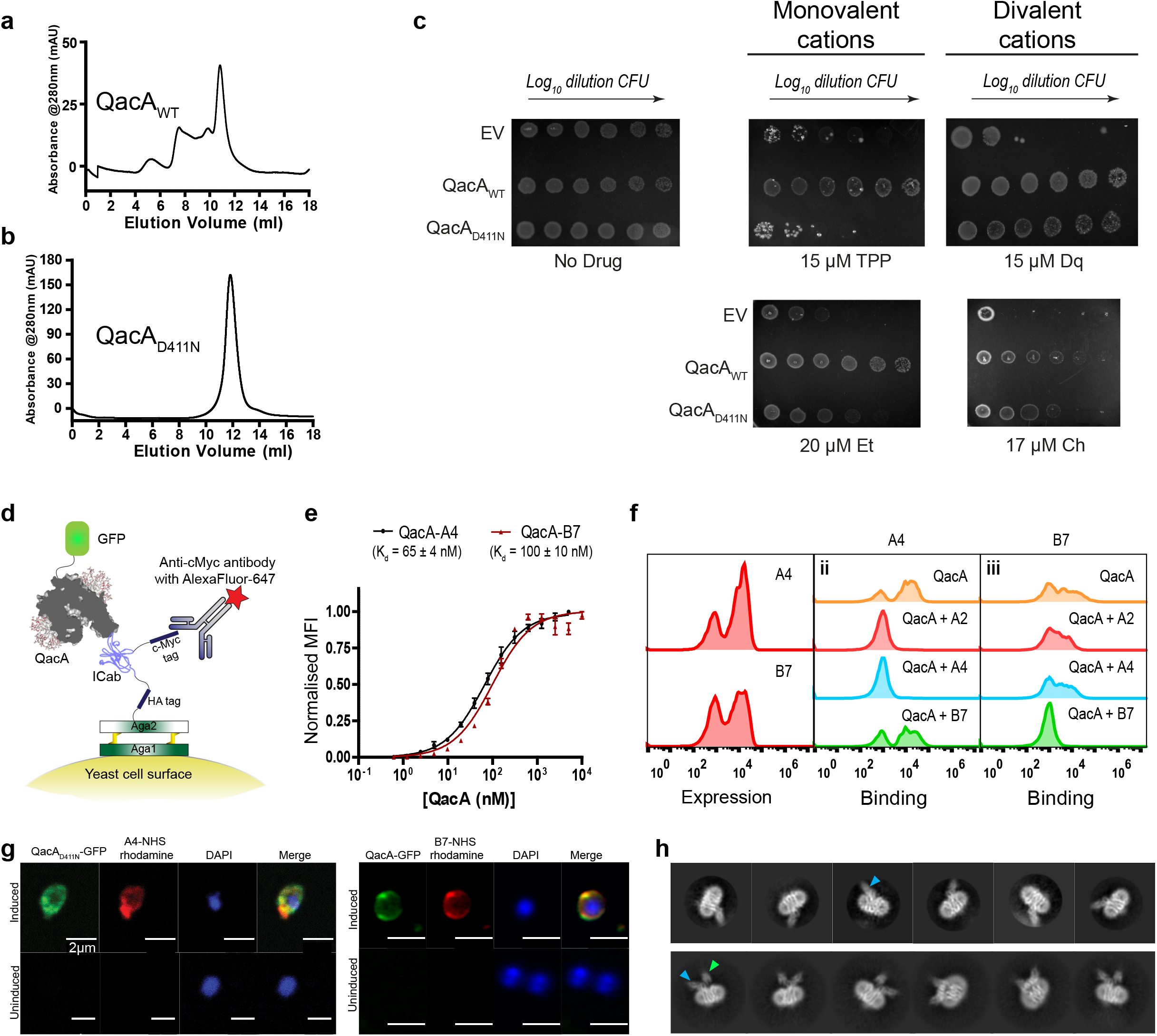
Generation of ICabs against QacA and their characterization. **a and b**, Size exclusion chromatography profiles of QacA_WT_ and QacA_D411N_ mutant, where the latter showed a more homogenous profile than earlier. **c**, Survival assays against substrates of QacA showed loss of activity against monovalent cationic antibacterials (TPP: Tetraphenylphosphonium, Et: Ethidium), while still retaining partial activity against divalent cations (Dq: Dequalinium, Ch:Chlorhexidine). EV: pBAD empty vector. n=3 for biological replicates. **d**, Schematic of yeast surface display platform used to enrich QacA specific ICabs from the cDNA library generated from camels immunized against QacA_D411N_. QacA was tagged with GFP while ICab was expressed augmented with c-Myc tag, probed by AlexaFluor-647 conjugated antibody. **e**, Population shift based titration of QacA-GFP against ICabs yield a *K*_d_ of 65 ± 4 nM for QacA-A4 complex and 100 ± 10 nM for QacA-B7. n = 3 for independent replicates. **f**, The yeast surface displayed A4 ICab could bind to pre-complexed purified ICab B7 and QacA-GFP and vice-versa, indicating that these ICabs do not compete for a common epitope and bind at different sites. Representational single set shown out of 3 independent replicates. **g**, Colocalization of QacA WT or QacA_D411N_ (tagged with GFP) on *E*.*coli* spheroplasts (stained with DAPI for viability) with ICabs (labelled with NHS-Rhodamine) was screened against ICab A4 (top panel) and ICab B7 (bottom panel). n=10-30 for number of spheroplasts imaged for each sample. **h**, Single particle CryoEM 2D class averages of QacA_D411N_ embedded in UDM micelle, in complex with ICab A4 (left panel) and with ICabs A4 and B7 (right), pointed with blue and green arrows respectively.

### Yeast display screening allows ICab isolation against QacA

Single-domain Indian camelid antibodies (ICabs) against QacA were screened through a yeast surface display platform using a previously optimized protocol (described in methods)^24^. ICabs were sorted for expression and affinity for QacA through FACS analyses (Fig. 1d, Supplementary Fig. 2a). Among the pool of binders, we found five unique clones that could induce population shifts in the range of 65 to 97% upon interacting with QacA_D411N_GFP (Supplementary Fig. 2b, c). Further analysis through QacA_D411N_ titration in flow cytometry yielded dissociation constants of 59 nM and 65 nM for ICabs A2 and A4, respectively. At the same time, ICabs B1, B2, and B7 displayed relatively weaker affinities in the range of 86-100 nM (Fig. 1e, Supplementary Fig. 3a).

We also evaluated whether the ICabs competed for a common epitope on QacA, through flow cytometry analyses. QacA_D411N_GFP was first incubated with a purified ICab and then screened for binding against yeast cells expressing other ICabs. In a situation involving a common epitope between the purified antibody and the competing antibody expressed on the yeast cell surface, we expect no shifts in the population as the purified antibody prevents interactions by the latter. In the scenario where a purified antibody binds to an alternate site that is distinct from the binding site of the surface-displayed antibody, the antigen (QacA_D411N_GFP) would interact despite the presence of the bound antibody. Through these analyses, we could establish that four among the five ICabs, including A2, A4, B1 and B2, interact competitively at the same epitope, whereas B7 shifts the population of cells despite having a bound antibody, suggesting interactions at an alternate epitope in a non-competitive fashion (Fig. 1f, Supplementary Fig. 3b). Similar results were obtained when ICabs A4 and B7 were heterologously purified, incubated with QacA_D411N_, and analyzed on FSEC (Supplementary Fig. 3c, d). These ICabs were also analyzed for topological localization using spheroplasts of *E*.*coli* expressing QacA-GFP, where we observed that both A4 and B7 bind to the extracellular surface of QacA (Fig. 1g). We employed these ICabs to perform electron cryomicroscopy (cryoEM) on QacA_D411N_ complexed with A4 and B7 with a combined mass of ∼80 kDa, excluding the detergent micelle. The reference-free 2D classes computed with the heterodimeric A4-QacA complex and the heterotrimeric A4/B7-QacA complex indicate their interactions on the extracellular face of QacA with independent albeit spatially proximal epitopes (Fig. 1h).

### CryoEM structure of QacA reveals outward-open conformation

The cryoEM structure of QacA_D411N_ in UDM micelles was determined in complex with two ICabs (A4 and B7) that served as fiducial markers during cryoEM image processing. The 2D classes of the particle set used for the final reconstruction displayed the presence of secondary structural elements within the transporter, indicating well-aligned particles with multiple orientations. The models of QacA and two ICabs were built into the coulombic map that was density modiInteraction of motif B arginine with the substrate fied (Supplementary Fig. 4, Supplementary table 1) and yielded unambiguous densities for nearly the entire transporter, from residues 5 to 514, and includes densities for all the 14 TM helices, and intra- and extracellular loops (Fig. 2a, Supplementary Fig. 5). ICabs A4 and B7 interact closely with the extracellular loop 7 (described later). Interestingly the QacA structure built into the cryoEM map tallies well with the outward-open structure of QacA’s AlphaFold2 model with a Cα rmsd of 1.3 Å for the entire molecule. The organization of the TM helices is typical of an MFS fold transporter where two six-helical bundles (TMs 1-6 and TMs 9-14) are arranged with a pseudo two-fold symmetry (Fig. 2a, b and c). The TMs 7 and 8 are arranged as an insert in the long loop joining the two bundles, and are juxtaposed next to the gap between TM2 and TM13 of QacA (Fig. 2c). This arrangement of TM helices is also observed in other 14 TM containing MFS transporters, including proton-dependent oligopeptide symporters (POTs) and NorC, a putative DHA2 member^24,25^. While the organization of helices is generally consistent with the MFS fold, the helical positions in QacA differ in comparison to the structures of DHA1 members like MdfA and LmrP (Fig. 2d). The differences are particularly apparent in the organization of TMs 2, 5, 9, 10, 13 and 14. The architecture of the QacA_D411N_ structure is outward-open with solvent accessibility clearly towards the extracellular side. The solvent-accessible vestibule is narrow compared to other DHA1 members, likely due to the discrepancies within helices that are placed relatively closer to the vestibule than LmrP and MdfA (Fig. 2e). The interdomain cleft between TMs 5 and 8 in LmrP and MdfA is wide open towards the extracellular side with relative angles of 41° and 45°, respectively, that can allow the interactions of amphipathic molecules like detergents through the bilayer (Fig. 2f)^26^. This cleft is narrow in QacA between TMs 5 and 10 (equivalent to TM8 in DHA1) as both helices are relatively parallel, disallowing the membrane environment to access the vestibule. The cleft formed between TMs 2 and 13 (TM11 in DHA1) widens towards the extracellular face of the transporter. However, exposure of the vestibule to the lipid bilayer in QacA is occluded by the presence of TMs 7 and 8 that are juxtaposed next to this cleft (Fig. 2g), whereas LmrP and MdfA lack this helical insertion^27^.

**Fig. 2.**
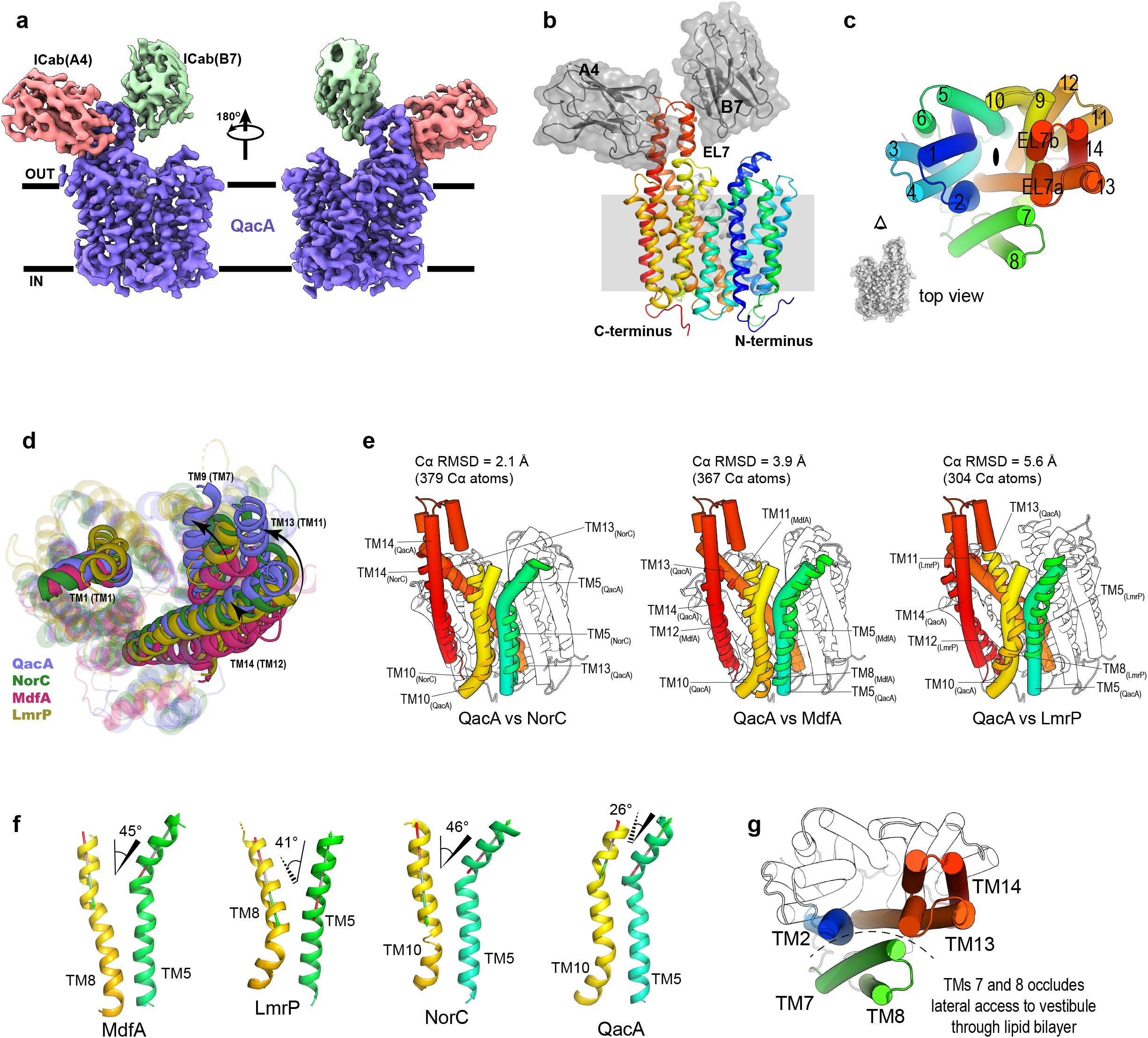
Architecture of QacA-A4-B7 complex. **a**, CryoEM model of QacA (blue) in complex with ICab A4 (pink) and ICab B7 (green). **b**, Cartoon representation of the complex, with 14 transmembrane helices of QacA shown in rainbow gradient from N-terminus to C-terminus. Extracellular loop 7 (EL7), with which B7 interacts, lies between elongated TMs 13 and 14 that protrude towards the periplasmic side of the membrane and form the epitope which interacts with A4. **c**, Topology of transmembrane helical arrangement, the top view shows canonical MFS fold of the transporter, along with the location of EL7 segment as seen in the top view. **d**, Structural alignment of QacA_D411N_ (blue) with MdfA (pink, PDBID: 6GV1), LmrP (yellow, PDBID: 6T1Z) and NorC (green, PDBID: 7D5P). Some of the major rearrangements in TM assembly are shown as opaque helices (TMs 9, 13 and 14 in DHA2 transporters QacA and NorC) with their corresponding TMs in DHA1 transporters MdfA and LmrP are labelled in parentheses. TM1 is shown to suggest a stationary point of view for these rearrangements. **e**, Side views of Structural alignments of QacA (cylindrical helices) with LmrP, MdfA and NorC with Cα RMSDs written above the alignments. TMs 5 (green), 10 (yellow, TM 8 in MdfA and LmrP), 13 (orange, TM 11 in MdfA and LmrP) and 14 (red, TM 12 in MdfA and LmrP) are highlighted to show structural differences between QacA and representative DHA transporters. EL7 of QacA is also shown as cylindrical helices. **f**, Angular separation between the upper halves of TM 5 and TM 10 in QacA and NorC, and of TM5 and TM8 in MdfA and LmrP. Dashed lines represent helices going into the plane to the tapered end, while solid line represents helices on the plane (uniform line) or coming out of the plane towards the broader end. **g**, cylindrical helices representation of QacA from top view to highlight block of lateral access of the vestibule to the lipid bilayer (dashed arc) due to presence of TMs 7 and 8 (green). TMs 2, 13 and 14 are shown in their usual colour scheme.

### The vestibule of QacA is negatively charged

The solvent-accessible vestibule in QacA extends halfway across the bilayer. It is secluded from access to the cytosol through multiple hydrophobic and H-bond interactions in TM helices that prevent solvent access towards the cytosolic half of the transporter below residue Trp149 in TM5, that forms the base of the vestibule (Fig. 3a). Like many other transporters of the DHA family, QacA has a negatively charged environment in its vestibule that facilitates the binding and transport of cationic antibacterials. However, as described in previous studies, unlike some well-characterized members of the DHA1 family, like NorA, MdfA and LmrP that use a single or a couple of acidic residues to drive transport, QacA possesses multiple acidic residues scattered throughout the vestibule, making the overall electrostatic potential inside the vestibule highly negative (Fig. 3a). These acidic residues have been categorized into three groups based on their contributions to substrate binding and transport^21^. While D34 and E407 have been deemed essential to QacA’s function for all types of substrates, residues like D411 and D323 are shown to play conditional roles in transporting specific divalent cationic substrates with longer linkers^21^.

**Fig. 3.**
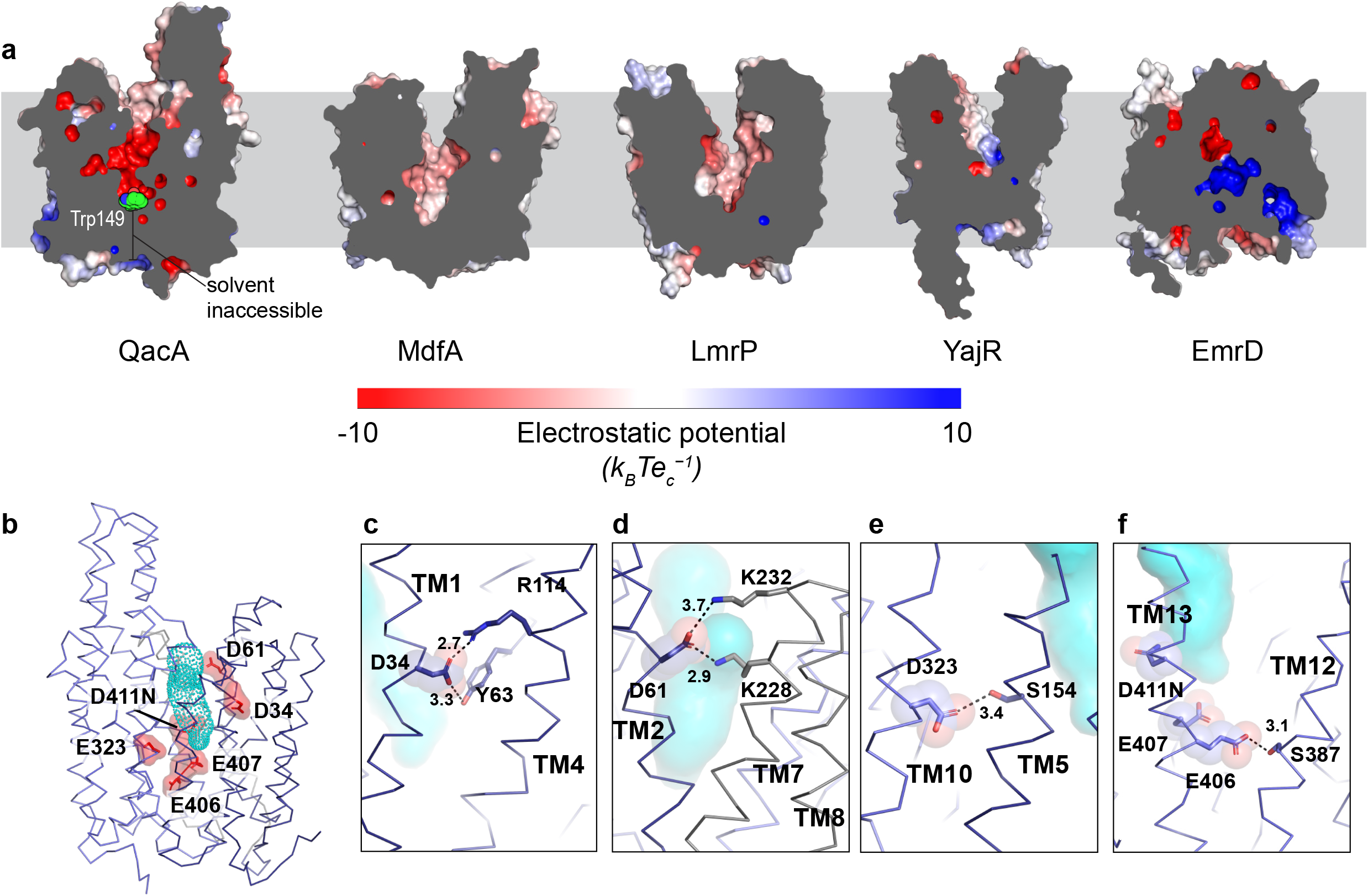
Solvent accessible vestibule in the outward open state of QacA_D411N_. **a**, APBS electrostatic maps in solid longitudinal sections of QacA along with common drug proton antiporters. Trp149 in QacA is shown in green spheres. **b**, Solvent accessible vestibule of QacA in this model (shown with blue dots) is lined with acidic residues, shown in red sticks and spheres. **c-f**, Neighborhood of all the acidic residues in the vestibule (sticks and transparent spheres; side chains of interactants are shown in sticks) with distances shown in angstroms (Å). Solvent accessible volumes are shown with cyan blobs in the background.

On the other hand, while providing partial fitness to transport substrates like ethidium and TPP, residues like D61 are considered non-essential for multiple substrates. E407 has also been attributed to having a moonlighting role of substrate recognition in the case of dequalinium while canonically acting like a protonation site during the transport of TPP and pentamidine. A homology-based inward-open model was earlier used to provide possible explanations for substrate binding and transport, and cysteine crosslinking-based assays of QacA mutants were used to verify relative solvent accessibilities of these residues^21^. The QacA structure described in our present study provides insights into the environment of the protonation sites in its outward-open state (Fig. 3b).

The acidic residues in the vestibule of QacA are involved in myriad polar interactions with neighboring residues through their side chains. D34 (TM1) interacts with a nearby R114 (TM4), and Y63 (TM2) (Fig. 3c). Y63 is conserved across transporters of the DHA2 subfamily (Supplementary Fig. 6) and is essential in the functioning of QacA, where a mutation to valine leads to loss of function in QacA but retains function upon Y63F mutation^28^. This type of interaction is also structurally conserved in the case of MdfA, a DHA1 transporter, where E26 interacts with Y30 (TM1) and Y127 (TM4), albeit only in the outward open conformation (PDB ID: 6GV1). Further, the D34 interacts with R114, a highly conserved residue that forms motif B in DHA transporters (Supplementary Fig. 6). Interaction of motif B arginine with the substrate recognizing acidic residues is suggested to enhance the pKa value and facilitate deprotonation at neutral pH^29^. On the other hand, D61(TM2), structurally very similar in its positioning along the normal to the lipid bilayer and with substantial solvent accessibility, has only partial conservation amongst DHA2 transporters but forms networks well within the vestibule with lysine residues from TM7 (K228 and K232) (Fig. 3d, Supplementary Fig. 6). This feature is absent in DHA1 transporters due to the lack of corresponding interdomain linker TMs 7 and 8 in them (Supplementary Fig. 6).

While there is no sequence conservation of D323 (TM10) even amongst DHA1 transporters, it fits within the groove formed by TM5 and TM10 between the N-terminal and C-terminal domains and forms a hydrogen bond with the hydroxyl group of Ser154 (TM5) (Fig. 3e). D323 is selectively needed to transport certain divalent cations like pentamidine, but not others^8,21^. Lying towards the lower leaflet along the normal of the bilayer, E406 and E407 in TM13 are peculiar in their partial conservation as acidic residues or as permanent protonation mimics as asparagines in many DHA2 transporters, which are sequentially similar to QacA. This is interesting, considering that E407 has been characterized primarily as a protonation site for most of QacA’s substrates, it acts as a substrate recognition site for some of QacA’s substrates like dequalinium and chlorhexidine that have cationic moieties separated by long linkers. E407 is exposed to the solvent-accessible vestibule, in QacA and could undergo protonation and deprotonation events during the transport cycle. E406 interacts with the hydroxyl group of S387 (TM12) through a hydrogen bond through its side chain (Fig. 3f). Comparing these residues with the corresponding ones from homologous transporters of the DHA2 family, one can find that many of these residues are conserved as such, or as their amide counterparts (Supplementary Fig. 6). However, as also seen with their surface electrostatics (Fig. 3a), the presence of multiple acidic residues is a phenomenon observed only in a certain subset of transporters like QacA, that also tend to harbor a conserved structured region in their extracellular loop 7 (EL7).

### EL7 organized as a hairpin loop is the epitope for ICab interactions

Unlike other DHA1 and DHA2 transporters, QacA TM helices 13 and 14 extend beyond the bilayer to form a highly ordered EL7. A comparison of the QacA and NorC structures displays the EL7 as a feature unique to QacA (Fig. 4a). A comparison of other canonical DHA2 members related to QacA display the presence of this insertion suggesting that EL7 hairpin loop is a distinctive structural feature (Fig. 4b, Supplementary Fig. 7). The residues in the region of 441 to 467 forms a helix-turn-helix motif that resembles a helical hairpin (EL7a, 7b) that is directed towards the membrane bilayer but does not occlude the vestibule of QacA in the outward-open conformation. It is notable that the EL7 region is characteristic of only a few transporters from the DHA2 family, but the region is structurally conserved, as seen in the AlphaFold2 models of SmvA and LfrA (Fig. 4c). The EL7 is the primary site of interaction with two ICabs albeit at different locations.

**Fig. 4.**
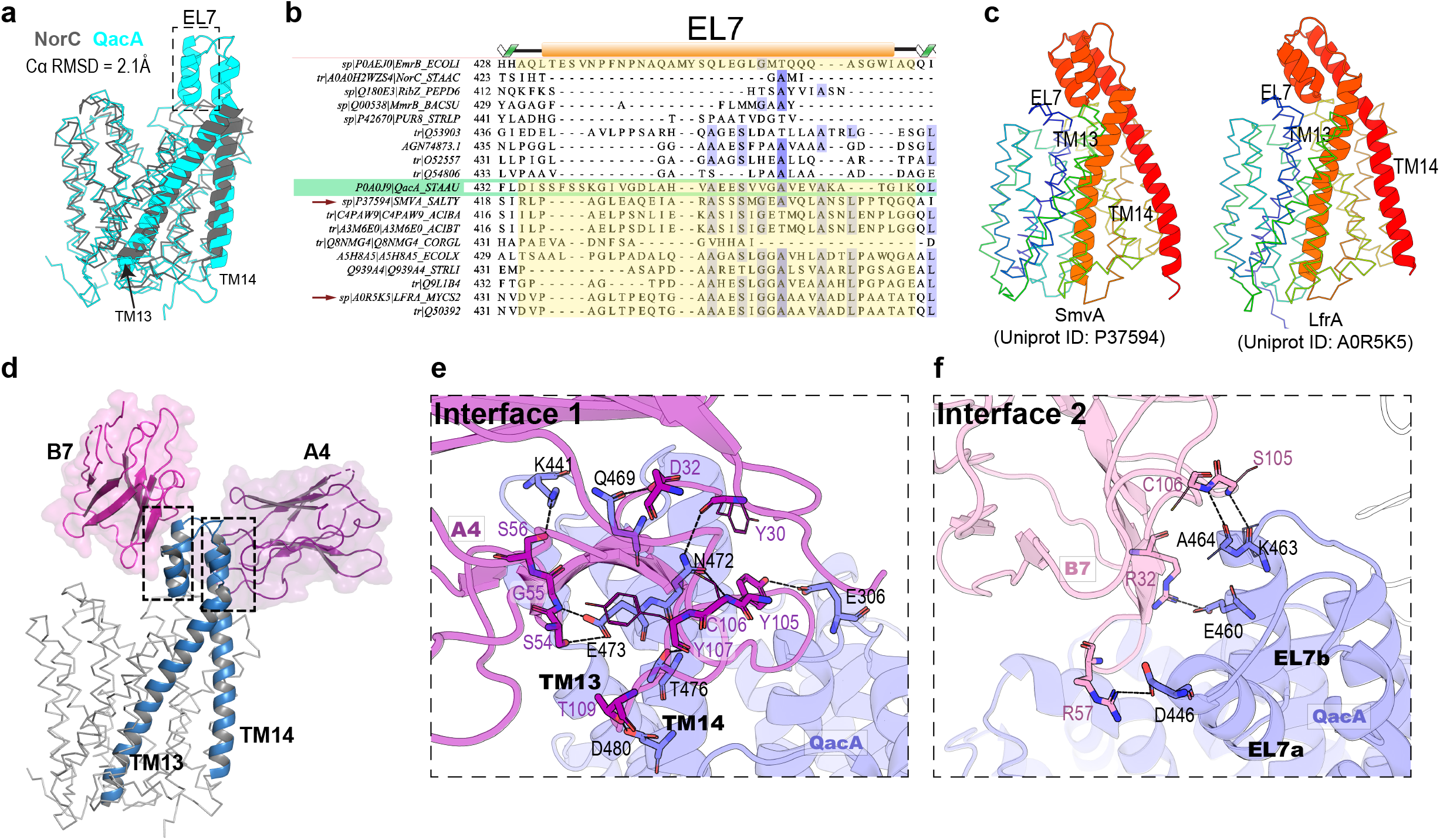
General overview of EL7 region of QacA, and QacA-ICab interfaces. **a**, Structural alignment of QacA_D411N_ (cyan, this study) and NorC (grey, PDBID: 7D5P) to depict absence of a structured EL7 in NorC. **b**, ROI of EL7 in Multiple sequence alignment of QacA and its homologs from various prokaryotic genera (Entire MSA in Extended Data Fig. 6). Stretches in proteins that possess EL7 are highlighted in yellow. QacA sequence is boxed in green. **c**, Ribbon representation of AlphaFold models of SmvA (*S. typhimurium*, Uniprot ID: P37594) and LfrA (*M. smegmatis*, Uniprot ID: A0R5K5) with TMs 13 and 14 and EL7 highlighted in cartoon, correlating with multiple sequence alignment as shown in **b. d**, Ribbon representation of QacA with TMs 13 and 14 and EL7 shown as blue cartoon. A4 and B7 nanobodies are shown in magenta and purple cartoons and surface respectively, with the epitopes highlighted as boxed ROI. **e**, Interface 1 (QacA-A4) and **f**, interface 2 (QacA-B7) shown with participating residues in pink and blue stick representations for ICabs and QacA respectively. Side chains of the residues which are interacting only through their main chain atoms are shown in thin lines for clarity.

The two ICabs interact at distinct sites of EL7 with A4 interacting with helical extensions of TM13 and TM14, and B7 interacting with the hairpin loop of QacA and holding it in place(Fig. 4d-f). The densities of the two ICabs are at a low resolution in some parts of the framework region but the antigen binding loops have well-defined densities that allowed chain tracing and side-chain assignment. ICab A4 displays a high affinity (*K*_d_=∼60 nM) for QacA EL7. This is the primary site where most of the binders interact. The ICab interacts with an interfacial area of 814.2 Å^2^. The epitope interactions are mediated through CDRs 1, 2 and 3, with CDR3 forming a bulk of the interactions and bridging the residues in EL6 (H362 and P363, VdW interactions) and EL5 (E306, H-bond with Y105 of CDR3 in ICab A4) with EL7 (Fig. 4e). The CDR3 of A4 is positioned unconventionally as it does not curve over the framework like a typical nanobody but elongates to form the antigen-binding site. CDRs1 and 2 have localized interactions with TM13 and TM14 extensions that form EL7. The ICab B7 is a conventionally structured camelid antibody and interacts weakly with the hairpin loop. The affinity displayed by B7 is weaker (*K*_d_=100nM) compared to A4 with also poorer density in the framework region leading to gaps in the residue assignment in the structure. The weakened affinity could be a consequence of a much smaller domain interface of 446.3 Å^2^. The interactions are mainly modulated through H-bonds mediated between EL7 hairpin and the residues in CDR loops 1 and 3 (Fig. 4f).

### Modeling alternate conformations of QacA

To delve into the mechanism of transport cycle in QacA, we predicted the inward-open state, QacA_io_, using AlphaFold2 that is known to predict transporter structures with high accuracy^30,31^. We also analyzed the range of motions in its transmembrane regions starting from outward-open state through equilibrium MD simulations, which helped us elucidate a cluster of micro conformations in the inward-facing occluded state represented by QacA_occ_. The Cα register and the secondary structure limits of QacA_io_ fit closely with the outward-open structure of QacA_D411N_ (Fig. 5a). To validate the QacA_io_ model further, we aligned QacA_D411N_ with both QacA_occ_ and QacA_io_, and compared vectors for each Cα displacement in the two alignment pairs. It should be noted that the displacements of residues towards the extracellular leaflet are necessary to occlude the vestibule from outer end, while those towards the intracellular leaflet would open the vestibule to solvent access from the cytoplasmic side, rendering the molecule inward-open. We found that most of the Cα displacements in both QacA_D411N_-QacA_occ_ and QacA_D411N_-QacA_io_ alignment pairs were similar in direction, with near identical shifts in Cα of EL7 and the residues lying towards the extracellular leaflet of TMs of N-terminal domain. This gave us greater confidence to derive predictions on the basis of tertiary structure level information derived from QacA_io_ model.

**Fig. 5.**
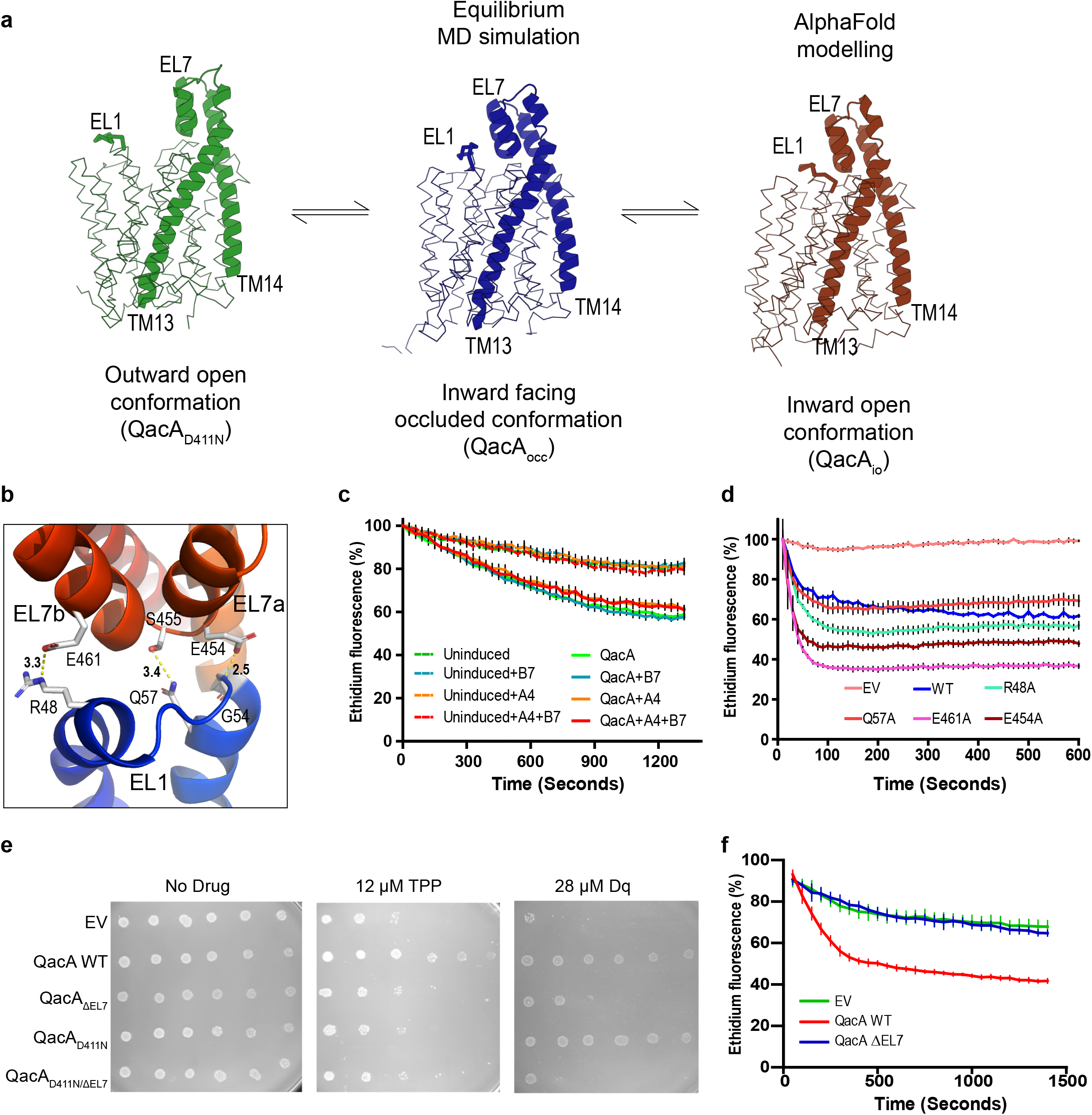
Structural overview of QacA-ICab complex interfaces. **a**, Ribbon representation of QacA with TMs 13 and 14 and EL7 highlighted in cartoon. Green, brown and blue representations are QacA_D411N_, QacA_io_ and QacA_occ_ states respectively. **b**, Apparent interaction network between residues of EL7 and EL1, as seen in QacA_io_ model. Distances between polar interactions are depicted and annotated in Angstroms. **c**, *E*.*coli* spheroplasts based efflux assay to assess the effect of nanobodies on ethidium efflux rate in QacA WT. n=6 for technical replicates; error bars represent SEM. **d**, Whole cell ethidium efflux assay with alanine mutants of interactants at EL1-EL7 interface. EV = empty vector. **e**, Survival assay with EL7 deletion construct (QacA Δ451-467) with QacA_WT_ or QacA_D411N_ in the background in TPP and dequalinium. **f**, Whole cell ethidium efflux assay showing loss of activity in QacA_ΔEL7_ mutant. n=3 for biological replicates, each in technical triplicates. Representative biological replicate with technical triplicates shown, with SEM marked with error bars.

### Structural organization and role of EL7

The conformational trajectory from QacA_D411N_ to QacA_io_ through QacA_occ_ reveals appearance of several interactions between residues of EL7 and extracellular loop 1 (EL1) present between TMs 1 and 2 in the N-terminal domain (Fig. 5b). While not all the interacting residue pairs are common to both QacA_occ_ and QacA_io_, we decided to check whether these pairs from the latter model had any role in regulating transport activity in QacA. In QacA_D411N_, B7 interacts with a patch on EL7 that should get occluded to free solvent upon its interaction with EL1, while A4 interacts with regions in TM13 and TM14 that are solvent exposed in both QacA_D411N_ and QacA_io_. It was reasonable that binding of B7 or A4 or both of these ICabs on QacA may interfere with this interaction between EL1 and EL7, thereby transport. To check for the same, we employed the use of *E*.*coli* spheroplasts to perform ethidium efflux assay in the presence of ∼10 times molar excess of ICabs. Since the electron transport chain is already present on the inner membrane of the cell, we proposed that the system will work in spheroplasts too. Indeed, we could see a steady efflux of ethidium from the spheroplasts with overexpressed QacA in the membrane when glucose was introduced in the medium, albeit more than an order of magnitude slower than when tried with whole cells. This could be attributed to the absence of a compartmentalized periplasm present in whole cells that helps in quickly re-establishing a pH gradient across the inner cell membrane. However, the rate of efflux did not change in the presence of B7, while reducing insignificantly in the presence of A4 nanobody, suggesting that they do not affect the transport of substrates through QacA in these concentrations (Fig. 5c).

Alanine mutagenesis of one or both residues in each of the interacting pairs was followed by functional assessment through whole cell ethidium efflux assay. We found that while the activities did vary to minor extents, we could not find any significant change in the transport activities of any of the EL7 alanine mutants of QacA, suggesting that single residue substitutions of interacting residues between EL1 and EL7 do not affect the transport activity in QacA (Fig. 5d). However, we see peculiar results when we delete the residues 451 to 467 that lie in the EL7 region of QacA. Upon EL7 deletion, even with comparative expression levels between QacA_WT_ and QacA_ΔEL7_, the latter fails to provide resistance against any of the antibacterial cations tested, as also reflected in the transport activity against Et in whole cell efflux assays, suggesting that even though disrupting interactions between EL1 and EL7 does not affect QacA’s transport activity, EL7 in its entirety may have a structural role in the conformational cycle of QacA (Fig. 5e, f).

### Asymmetric rocker-switch movements in QacA

Superposition of the three conformations of QacA reveals an unconventional pattern of alternating-access in QacA during its transport cycle. A zoomed-out view of the superposition also gives us a rough center of rotation about which the five TMs move in the N-terminal domain, lying near the middle of the bilayer. In the N-terminal domain, individual transmembrane helices TMs 1-2 and TMs 4-6 undergo angular motions ranging between 17° and 30° about a point lying roughly at the center of the bilayer (Fig. 6a). However, such significant motions are not seen in the C-terminal domain, except in EL7, which bends inwards towards the vestibule, in QacA_io_ (Fig.6b, c). This result is not an artifact of structural alignment as we also analyzed local motions within the domains of QacA by calculating distances between every Cα pair within a conformation, and checked if any of these distances changed between the two states QacA_D411N_ and QacA_io_ (Supplementary Fig. 7). We performed the same analysis for two more transporters MdfA and LacY, where both are members of the major facilitator superfamily, but the earlier is a DHA1 member, while the latter is a proton-dependent sugar symporter. These transporters have been structurally elucidated in both outward and inward open conformations, providing us with reference conformations to compare our data with. For a pair of Cα atoms in the same domain, if the distance between them changes, we infer that the domain undergoes local motion between those residues. As predicted and shown in Supplementary Fig. 7, we see larger local motions between residues of the N-terminal domain than in C-terminal domain in QacA, concurring with our previous observations (Fig. 6b-d).

**Fig. 6.**
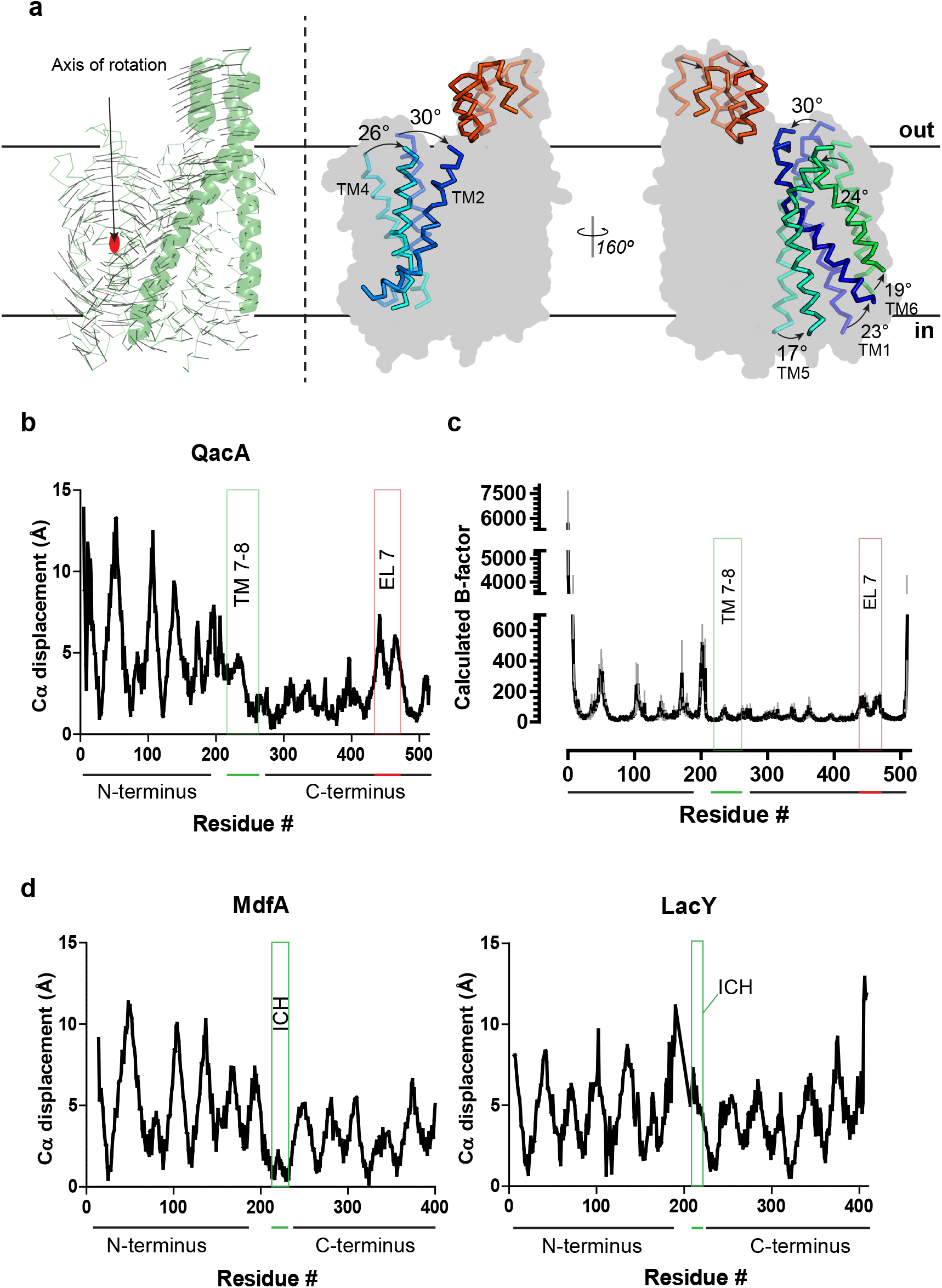
Asymmetric rocker switch mechanism of transport in QacA. **a, (Left)** The Cα displacement vectors between aligned QacA_D411N_ structure and QacA_io_ model showing axis of rotation traversing along and through the middle of the bilayer to depict asymmetry between the degree of motions between N-terminal and C-terminal domains. TMs 13-14 and EL7 are shown as cartoon helices. **(right)** Major TM displacements seen between QacA_D411N_ and QacA_io_ models are shown with a background of QacA_D411N_ boundary to depict their locations in the transporter. Faded ribbons are from QacA_D411N_ structure, while opaque ribbons are from QacA_io_ model. Average angular displacements for each TM region are marked. Note that some helices are labelled with two different angular motions around a point as different regions in the TM undergo different extents of motions. **b**, Cα displacement graphs for QacA, with extents of N- and C-terminal domains, TMs 7-8 and EL7 annotated. **c**, RMSF plot for MD simulations with QacA_WT_ modelled on QacA_D411N_ structure as the initial state show good correlation with Cα displacement plot for QacA in **c** panel. RMSF values shown as calculated B-factors using the conversion (800π^2^(RMSF)^2/3^). Error bars show SEM of data from n=2 for independent replicates. **d**, Cα displacement graphs for MdfA (left) and LmrP (right), with extents of N- and C-terminal domains and intracellular helical segment (ICH) annotated.

We conclude that although a certain extent of asymmetry in rocker-switch motion exists in all the three MFS transporters analyzed here, the asymmetry is much more prominent in case of QacA, where the TMs 1, 2, 4, 5 and 6 of the N-terminal domain move to a greater extent than the TMs of the C-terminal domain. The motions in the TMs are complemented with the extracellular loops EL1 and EL7, which form a gate as QacA shifts from outward open conformation to inward open. While these interactions do not have a regulatory effect of QacA’s transport activity, the EL7 does have a structural role to play in QacA’s conformational landscape. Given the similarity of structural organization in models of DHA2 members like SmvA and LfrA with QacA, the asymmetric rocker-switch motion could be extended to these transporters as well.

## Discussion

The study delves into the structural organization of a prototypical member of DHA2 transporters, QacA, in complex with two ICabs that do not interfere with QacA’s efflux activity. The structure of QacA displays the symmetric organization of two helical bundles 1-6 and 9-14 with the TMs 7 and 8 forming an inserted helical pair (6+2+6 arrangement) characteristic of 14TM helix MFS transporters. The structure of QacA represents an outward-open conformation with solvent access visible to nearly halfway across the membrane bilayer. In comparison to the better-studied DHA1 members, QacA displays a significantly larger number of acidic residues that surround the vestibule and play distinct roles, ranging from being essential for substrate recognition and acting as protonation sites for selected substrates, to having merely structural roles without a direct bearing on cation interactions. The structure elucidation of QacA provides the precise locations of these acidic residues, and allows visualization of their environment and interactions within the transporter.

The distinct structural feature of QacA is the EL7 that is organized as a hairpin loop inserted between TMs13 and 14. This is a unique insertion that is observed in multiple DHA2 members like LfrA and SmvA whose structural context was previously unknown. The presence of this EL7 hairpin makes QacA and its related sequences, a distinct subset among the DHA2 transporters. The region of TM13-EL7 hairpin-TM14 helical bundle that is solvent exposed is also the site of interactions for the two ICabs used in the study. Unlike recent studies on NorA and NorC where the antibodies interact by inserting the CDR loops into the vestibule of the transporter and lock the transporter into an outward-open state^13,22^, the ICabs interact with EL7 at distinct locations and hold it in place resembling a “molecular bow-tie”. Surprisingly, despite the interactions with two ICabs, the efflux activity of QacA, measured in spheroplasts, remains unhampered. While this is an unexpected result, it is not out of place in some contexts observed earlier, for instance in LacY, diverse nanobodies do allow substrate interactions and affect structural dynamics of LacY to varying extents^32^.

Comparison of the outward-open state with an occluded conformation of QacA obtained through molecular simulations and AlphaFold2 model of QacA in the cytosol-facing conformation reveal the formation of an interface between EL7 hairpin and EL1. While alanine substitutions at the interface residues did not interfere with the transport properties, the deletion of the hairpin loop did severely compromise the efflux properties of QacA indicating the important structural role EL7 plays in DHA2 members.

We further observe using these models that the helical bundle of TM1-6 undergoes significantly larger movements in comparison to the TMs 9-14 that remain relatively static and play a scaffolding role. The centre-of-mass of the TM helices 1-6 remains unaltered in its position during these shifts indicating the rotation of the helical bundle around a single axis without any translation associated with it. This is in stark contrast with the other MFS members like LacY and the DHA1 member MdfA whose structures are available in both outward and cytosol-facing states^14,27,33,34^. Although the study does not have an experimental QacA structure facing the cytosol, the accuracy of AlphaFold2 models and the simulations done with QacA outward-open coordinates provide the insights to suggest the presence of an “asymmetric rocker-switch” in QacA and other members of DHA2 family that have a similar structural organization.

The study provides the template for a detailed understanding of QacA’s interactions with antibacterial compounds and its conformational shifts to mediate proton-coupled efflux in *Staphylococcus aureus* that would aid in the design of efflux pump inhibitors against QacA and other DHA2 members.

## Acknowledgements

Research in the manuscript was supported in part by the Wellcome Trust/DBT India Alliance Intermediate Fellowship (IA/1/15/2/502063) awarded to AP. Research support through Dept. of Biotechnology, India (DBT) grant (BT/PR31976/Med/29/1421/2019) to AP is acknowledged. AP is an EMBO Global Investigator from India. PM, SA, PA, AA are graduate students funded by the Indian Institute of Science. RR is a principal scientist at the NRCC, Bikaner. We acknowledge Diamond Light Source for access and support of the cryo-EM facilities at the UK’s national Electron Bio-imaging Centre (eBIC) [proposal EM BI29692-1], funded by the Wellcome Trust, MRC and BBRSC. The authors thank Dr. Davide Zabeo for data collection at EBIC. We thank Vinothkumar KR and Sucharita Bose and the National Electron Cryo-Microscopy facility, BLiSc, Bangalore (DBT/PR12422/MED/31/287/2014) for initial screening preliminary data collection of the grids. We would like to acknowledge Somnath Dutta at the advanced cryoEM facility, IISc funded through DBT-BUILDER program (BT/INF/22/SP22844/2017) and DST-FIST program (SR/FST/LSII-039/2015) for screening and data collection. Computational support from the high-performance computing facility “Beagle” setup from grants by a partnership between the DBT, India and the Indian Institute of Science (IISc-DBT partnership programme) is acknowledged. The authors would like to acknowledge the DBT-IISc partnership program phase II and the DST-FIST program support. We thank Prof. Raghavan Varadarajan for access to FACS facility.

## Methods

### Plasmids and Strains

JD838 (Δ*mdfA*Δ*acrB*Δ*ydhE*::*kan*) strain derived from *E. coli* K-12 strain, LMG194, bearing a knockout of 3 multidrug efflux pumps^1^ was used for all the experiments in this study if not mentioned otherwise. A codon-optimized QacA synthetic gene^35^ was cloned into a pBAD-His_8_ vector between *NdeI* and *HindIII* sites. C-terminal GFP constructs were generated using the megaprimer based whole plasmid PCR method. Site-directed mutagenesis was performed to modify WT QacA to a stable functional construct QacA_D411N_ and QacA_D411N_GFP and cloned into pBAD-His_8_ vector. Extracellular loop (EL) deletion constructs QacA_ELΔ451-467_, QacA_ELΔ451-467_GFP, QacA_D411N-ELΔ451-467_, and QacA_D411N-EL/Δ451-467_ were cloned using the megaprimer based whole plasmid PCR method into pBAD-His_8_ vector.

For QacA construct optimization process, the cytosolic rim mutants E137Q, E141Q, R142Q, A143N, D267N, L389N and L392N were constructed using site-directed mutagenesis with specific primers and megaprimer based whole plasmid PCR was used to clone them in QacA_D411N_ pBAD-His_8_ backbone. Similarly, for EL1-EL7 interaction studies, R48A, Q57A, E461A and E454A mutants, and a deletion construct of Δ451-467 (ΔEL7) in the background of pBAD-QacAwt were generated. The protein expression profiles were checked using JD838 strain of *E. coli*. The ICab library was cloned into the pETCON vector between *NdeI* and *XhoI* restriction sites. The ICab library was expressed in *Saccharomyces cerevisiae* EBY100 strain, used for FACS and flow cytometry analysis. The ICabs were sub-cloned in pET22b vector without pelB signal sequence and over-expressed in SHuffle T7 cells of *E*.*coli* (NEB, catalogue no. C3026J).

### QacA protein purification

QacA and its mutants used in this study were purified following the protocol as described in our earlier study^21^. The purified membrane was homogenized in lysis buffer (30 mM phosphate buffer pH 7 and 120 mM NaCl, 5 % glycerol). The protein was extracted using 20 mM UDM detergent and kept for solubilization for 1 hour at 4ºC. The remaining cell debris and insoluble membrane were removed using ultracentrifugation at 30,000 rpm at 4ºC for 1 hour. The supernatant was kept for binding with pre-equilibrated Ni-NTA beads for 1 hour at 4ºC. The beads were washed with 30 mM imidazole containing wash buffer (30 mM phosphate buffer pH 7 and 120 mM NaCl, 5 % glycerol, 1 mM UDM) and eluted in the same buffer with 300 mM imidazole. The freshly eluted protein was quickly concentrated using a 50 kDa cut-off concentrator and purified further by size exclusion chromatography using Superdex 200 increase 10/300 GL column.

### Nanobodies library generation

Nanobodies library was generated as described. A male camel of age 4-5 years was immunized with purified QacA_D411N/E137Q_ protein in detergent micelle and reconstituted in proteoliposomes in prime and 7 boosters’ regime. The blood was collected after seven days of final immunization and PBMCs (peripheral blood mononuclear cells) were isolated. The total RNA was purified from PBMCs, and reverse transcribed into the cDNA using the Thermo scientific RevertAid First Strand cDNA Synthesis Kit. The VHH open reading frame was amplified from cDNA, and the PCR product along with the linearized pETCON plasmid was electroporated in EBY100 strain of *S. cerevisiae*. Positive transformants were selected and grown in SDCAA media, and induced in SGCAA media, followed by FACS sample preparation as described^36^. The cells were labelled with anti-HA antibody conjugated to Alexafluor 647 (catalogue no. 26183-A647), to estimate the levels of expression and QacA-GFP was used to estimate binding of the nanobodies on the yeast cell surface. The cells that were positive for both the expression and QacA-GFP binding were enriched for three rounds using FACS. A concentration of 1000 nM, 300 nM and 100 nM QacA-GFP was used in first, second, and third round of sorting respectively and twenty million cells were sorted in each round on BD Aria III cell sorter. Expression and binding of unique individual ICabs clones were analyzed on Aria C6 flow cytometer as described^36^. To estimate the expression of individual Icabs, anti-c-Myc antibody conjugated to Alexafluor 647 (catalogue no. MA1-980-A647) was used. Affinity of individual ICabs for QacA_D411N_ on the yeast cell surface was estimated using flowcytometry as described in earlier studies^24,37^.

### ICab expression and purification

ICabs (A2, A4, and B7) were cloned into pET22b vector and expressed in the SHuffle T7 *E*.*coli* cells. The cells were induced with 1 mM IPTG and grown at 30ºC. The harvested cells were resuspended in the lysis buffer (HEPES 30 mM pH 7.4, NaCl 120 mM) and sonicated. The cell debris and insoluble aggregates were removed using centrifugation at ∼18500*g* for 1 hour and the supernatant was kept for binding with pre-equilibrated Ni-NTA resin. The column was washed with 50 column volume of wash buffer (HEPES 30 mM pH 7.4, NaCl 120 mM, Imidazole 30 mM) and the protein was eluted with elution buffer (HEPES 30 mM pH 7.4, NaCl 120 mM, Imidazole 300 mM). The purified protein was further purified by size exclusion chromatography using Superdex 75 10/300 GL column.

### Epitope Competition assay with ICabs displayed on the yeast cell surface

To check the epitope competition between ICabs, QacA_D411N_GFP (100 nM) was incubated with purified ICab A2/ A4/ B7 (1000 nM) for 30 minutes at 4°C. This preformed complex of QacA_D411N_GFP-A2/ A4/ B7 was incubated with the yeast cells expressing ICabs on their cell surface. No binding was observed if the yeast cell surface displayed ICab and the purified ICab shared the same epitopic region suggesting competitive interactions.

### Spheroplasts preparation

pBAD-His_8_ QacA_D411N_GFP transformed *E. coli* JD838 cells were grown overnight at 37°C. QacA-GFP overexpression was induced in the secondary culture (5 ml)with 0.05 % L-arabinose and the cells were harvested as the OD_600_ reached 0.5-0.7. The harvested cells were washed with 1x PBS, 0.1 % BSA solution. The cell pellet was resuspended in 500 µl of 800 mM sucrose solution. In order to form the spheroplasts the following solutions were added in the described order: i) 30 µl of 1M Tris-HCl pH 8, ii) 24 µl of 0.5 mg/ml lysozyme, iii) 6 µl of 5mg/ml DNase, iv) 6 µl of 125 µl EDTA-NaOH pH 8 and v) 5 µl of 1 mM of spermidine. The mixture was incubated at room temperature for 20 minutes. The spheroplasts were harvested by low-speed centrifugation at 500g for 3 minutes. To stabilize the spheroplasts, 4 µl of 1 mM Tris-HCl pH 8, 400 µl of 800 mM sucrose, 8 µl of 1 mM MgCl_2,_ and 4 µl of 1 mM of spermidine were added to the spheroplasts and the volume was brought upto 400 µl with sterile water.

### Confocal sample preparation

ICabs A4 and B7 were labelled with NHS-Rhodamine (5/6-carboxy-tetramethyl-rhodamine succinimidyl ester, ThermoFisher Scientific (catalogue no. 46406)) as described by the manufacturer. For conjugation, a molar ratio at 10:1 for NHS-Rhodamine : protein was used in the conjugation buffer 20 mM HEPES pH 7. The reaction was stopped by the addition of Tris buffer pH 8. The non-reacted dye was removed using PD-25 Desalting Columns. The NHS-Rhodamine labelled ICabs were added to the freshly prepared spheroplasts at a final concentration of 0.03 mg/ml. The spheroplasts were incubated at room temperature for 30 minutes and washed thrice with 1x PBS, 0.1 % BSA solution to remove unreacted ICabs. Further, 10 ug/ml final concentration of DAPI was added to spheroplasts. The spheroplasts were washed once again to remove extra DAPI with 1x PBS, 0.1 % BSA solution. The spheroplasts were then fixed with a drop of ProLong™ Gold Antifade Mountant (Thermo Scientific, product code P10144). The slides were observed under confocal microscope (Olympus FV3000 or ZEISS LSM 800) for DAPI (λ_Ex_/λ_Em_ 358/461), NHS-rhodamine (λ_Ex_/λ_Em_ 552/575) and GFP (λ_Ex_/λ_Em_ 488/512).

### Binding study using FSEC

Purified QacA_D411N_ or QacA_D411N-ELΔ451-467_GFP was mixed with ICab (A4, B7) at a 1:1.2 ratio, and incubated at 4ºC for 1 hour. The shift in FSEC profile was observed by comparing QacA_D411N_-ICab complex and QacA_D411N_. The intrinsic tryptophan fluorescence was measured λ_Ex_ 294 nm and λ_Em_ 334 nm.

GFP-tagged crude membrane protein sample was prepared as described above. Purified ICab A4 or B7 were mixed with the solubilized QacA_D411N_GFP and QacA_D411N-ELΔ451-467_GFP samples and incubated at 4ºC for 1 hour. GFP-fluorescence was measured to compare the elution volumes of free QacA and QacA-ICabs complex..

### Cryo-EM sample preparation and data collection

SEC purified QacA_D411N_ in 120 mM NaCl, 2% glycerol and 1 mM UDM buffer was concentrated to 4 mg/ml using a 100 kDa cut-off centricon. QacA_D411N_ and ICabs A4 and B7 were mixed in a 1:1.2:1.2 (QacA:A4:B7) molar ratio. To ensure QacA_D411N_-A4-B7 complex formation, a small aliquot from concentrated peak fraction of QacA_D411N_-A4-B7 complex was analyzed using FSEC. The concentrated protein sample was ultracentrifuged at 30 k rpm at 4ºC for 1 hour just before the grid freezing. Quantifoil gold (R 0.6/1) gold holey carbon grids were used to freeze QacA_D411N_-A4-B7 protein sample using FEI Vitrobot. The grid freezing was carried out at 100% humidity, 5 s blot time, 10 s wait time, and 16ºC temperature, without blot force. Prelimainary data were collected on a Titan Krios equipped with a K2 detector followed by screening of optimized grids using Talos Arctica 200 keV cryo-electron microscope and the best grids were used for the final dataset collection. The final dataset was acquired on Titan Krios 300 keV electron microscope equipped with Gatan K3 detector at the EBIC-Diamond Light Source, UK (KriosI-m02). A total of 7770 movies were collected in super-resolution mode, 2x binning with a pixel size of 0.831 Å/pixel. The total electron dose was 51 e^-^/ Å^2^ with an exposure time of 2 s and an energy filter slit of 20 eV. The electron dose was fractionated over 50 frames.

### Cryo-EM data processing and refinement

The data was processed using cryoSPARC^38^. The imported 7770 movies were motion corrected using patch motion correction and CTF estimation was done using patch CTF estimation. The poor-quality micrographs were removed using a manually curate exposure tool present in cryoSPARC^38^ with a cutoff of 1.5 ice thickness and CTF correlation greater than 5.0 Å and a total of 6310 micrographs were selected for further processing. 158,327 particles were picked automatically using the blob picker tool from 400 micrographs and using inspect particle pick 136,503 particles were chosen. The 2D classification was performed with extracted 107,759 particles. Twelve 2D classes were selected, which consisted of 26,890 particles. These particles were used as templates for the template picker tool and a total of 5,723,316 particles were picked from 6310 micrographs. Particles were further inspected using inspect particle pick tool and 2,276,055 particles were selected. 1,666,340 particles were extracted from the selected particles, and 2D classification was performed. From the first round of 2D classification 502,364 particles were selected and further 2 rounds of 2D classifications were performed to remove noise. Finally, from the last 2D classes 218,040 particles were selected which comprises 97 2D classes. A single ab-initio model was built with 218,040 particles and homogeneous refinement was performed with a maximum align resolution of 3 Å and keeping other values as default. The GSFSC value after homogeneous refinement was 4.85 Å. Further, non-uniform refinement was performed taking previously refined volume as input. In the case of non-uniform refinement^39^, the number of extra final passes was changed to 5, maximum alignment resolution was changed to 2 Å and GSFSC fit resolution was kept as 5 Å. Also, the initial batch size was changed to 2000 with a batch epsilon value of 0.01 and the dynamic mask start resolution was kept as 7 Å. The value of GSFSC resolution after non-uniform refinement became 3.84 Å following the standard FSC cut-off value of 0.143. Further, the map resolution was significantly improved by density modification using the Phenix Resolve density modification tool^40^, and the final resolution after density modification was 3.6 Å. QacA_D411N_ and ICabs A4 and B7 AlphaFold2 models were preliminarily fitted into the map using Chimera^41^ and the model was built using Coot^42^. The model was refined using Phenix real-space refinement programme to remove the outliers and improve the model refinement statistics^43^. After iterative real space refinement, model matches to the density modified map with a CC of 0.77 (Supplementary Table1). The local resolution of the map was calculated using local refinement in Phenix^40^.

### Drug-resistance assay

Drug resistance assay was performed in the presence of dequalinium (Dq) and TPP with JD838 cells expressing WT QacA-GFP, WT QacA_ELΔ451-467_GFP, QacA_D411N_GFP, QacA_D411N-ELΔ451-467_GFP, and cells with empty pBAD vector. The experiment was performed in solid media 2 % w/v Luria broth and 1.8 % w/v agar. Cells were diluted to an OD_600_ of 1, and a series of 10-fold dilutions were prepared, and 2 µl from each dilution was spotted on Luria broth-agar plates. The plates contained 0.05 % (w/v) L-arabinose and 100 µg/ml Ampicillin, with or without the addition of toxic substrate Dq (18 µM, 24 µM, 28 µM) and TPP (12 µM, 15 µM). The survival of the cells was checked after 14 h of incubation at 37ºC to check the resistance against Dq and TPP.

The expression of WT QacA-GFP, QacA_ELΔ451-467_GFP, QacA_D411N_GFP and QacA_D411N-ELΔ451-467_GFP of the same set of cells was checked using FSEC.

### Whole cell and Spheroplasts based Ethidium efflux assay preparation

Whole cells-based ethidium efflux assay was done as previously described (ref JMB). JD838 cells harboring pBAD-*QacAwt* and its mutants, or empty vector were inoculated in 2% Luria Broth (w/v) which were used as 1% inoculum for 10ml of secondary cultures. The cells from the secondary culture were induced with 0.05% (w/v) L-arabinose and grown till the OD_600_ reached 0.6. The cells were washed with 20mM Hepes pH 7.0 twice, incubated with 20µM EtBr and 0.5 µM carbonyl cyanide m-chlorophenyl hydrazone (CCCP) for 1 hour in dark at room temperature. The cells were washed thrice and finally resuspended in 2ml of 20 mM Hepes pH 7.0 for the efflux assay.

For preparation for spheroplast based efflux assay, cultures were grown as described above for JD838 cells harboring pBAD-QacA-wt. Secondary culture grown in 100ml of 2% Luria Broth with 100 µg/ml ampicillin was split into two equal volumes. 0.1% w/v L-arabinose was added to one of the 50ml cultures to induce QacA over-expression, while the other was left uninduced. When the cultures’ OD_600_ reached 0.6 A.U., equal number of cells were harvested by centrifugation at 3000*g* for 15 minutes and washed twice with 1x Phosphate buffer saline (pH 7.4). The cells were incubated with 20uM EtBr for 10 minutes and then washed thrice with PBS. The cells were incubated in spheroplasting solution (0.5 µM CCCP, 700mM sucrose, 10 mM EDTA, 20mM Tris pH 7.2 and 0.2mg/ml lysozyme) for 20 minutes at 4°C with constant and slow tumbling. The spheroplasts were stabilized by adding 10 ml of ice-cold stabilizing solution (700mM sucrose, 20mM MgCl_2,_ and 20mM Tris pH 7.2) to every 30 ml of spheroplast suspension. The spheroplasts were harvested and washed thrice by centrifugation at 1000*g* for 15 minutes each and resuspended again in 5 ml of resuspension solution (650mM sucrose, 5mM Tris pH 7.2, and 10mM MgCl_2_).

### Ethidium efflux assay measurement

The assays were done in a 96 well corning flat bottom black plate with 90 µl of cells or spheroplasts. ICabs were purified in 10mM Tris pH 7.2 buffer and 150 mM NaCl for the spheroplasts based assay, and the same buffer was used as buffer control in the assay. A final concentration of 0.2 mg/ml of each ICab was used to incubate with spheroplasts. The cells were energized with 5mM whereas the spheroplasts were energized using 2% (w/v) of glucose and fluorescence measurements at λ_Ex/Em_ = 530nm/610nm were done on Thermo Varioskan Flash with 100ms long measurements after every 30 seconds for 30 minutes.

### Molecular Dynamics Simulations

A simulation box was generated for the coordinates of QacA_D411N_ where residue 411 was reverted to aspartic acid, using CHARMM-GUI^44-46^. CHARMM36m forcefield was used for the simulation run on Gromacs-2020 MD engine. All the acidic residues that were solvent exposed were kept deprotonated except D411, which was modelled with a neutral side chain. The set up contained >124000 atoms made of QacA, 224 and 70 molecules of POPE and POPG respectively in the bilayer, and charge neutralized with ∼150mM NaCl and water molecules. The system was energy minimized using steepest gradient descent and equilibrated step-wise with decreasing positional restraints. Berendsen and Nose-Hoover thermostat were used for temperature coupling, and Berendsen and Parrinello-Rahman barostats with sem-isotropic p-coupletype were used for pressure coupling, during equilibration and production MD runs respectively. After box preparation, energy minimization was done independently twice to generate two different random seeds as replicates. Simulations were also run for systems with a deprotonated D411 and N411 with identical parameters, but desired conformational changes were not observed for the 1µs simulations done for each of the simulations. More than 2µs of MD trajectory of neutral D411 QacA was used for analysis.

## Data Availability

Cryo-EM density map has been deposited in the Electron Microscopy Data Bank under accession number EMD-33612. Model coordinates have been deposited in the Protein Data Bank under accession number 7Y58. All other data needed to evaluate the conclusions in the paper are present in the paper and/or the supplementary materials. Source data will be provided with this paper upon acceptance of the manuscript.

## References

1. Blair, J.M., Webber, M.A., Baylay, A.J., Ogbolu, D.O. & Piddock, L.J. Molecular mechanisms of antibiotic resistance. Nat Rev Microbiol 13, 42–51 (2015).

2. Piddock, L.J. Multidrug-resistance efflux pumps - not just for resistance. Nat Rev Microbiol 4, 629–36 (2006).

3. Chitsaz, M. & Brown, M.H. The role played by drug efflux pumps in bacterial multidrug resistance. Essays Biochem 61, 127–139 (2017).

4. Pu, Y. et al. Enhanced Efflux Activity Facilitates Drug Tolerance in Dormant Bacterial Cells. Mol Cell 62, 284–294 (2016).

5. WHO. Antibiotic resistance: Multi-country public awareness survey I. World Health Organization. (2015).

6. Tong, S.Y., Davis, J.S., Eichenberger, E., Holland, T.L. & Fowler, V.G., Jr. Staphylococcus aureus infections: epidemiology, pathophysiology, clinical manifestations, and management. Clin Microbiol Rev 28, 603–61 (2015).

7. Costa, S.S., Viveiros, M., Amaral, L. & Couto, I. Multidrug Efflux Pumps in Staphylococcus aureus: an Update. Open Microbiol J 7, 59–71 (2013).

8. Paulsen, I.T., Brown, M.H., Littlejohn, T.G., Mitchell, B.A. & Skurray, R.A. Multidrug resistance proteins QacA and QacB from Staphylococcus aureus: membrane topology and identification of residues involved in substrate specificity. Proc Natl Acad Sci U S A 93, 3630–5 (1996).

9. Dashtbani-Roozbehani, A. & Brown, M.H. Efflux Pump Mediated Antimicrobial Resistance by Staphylococci in Health-Related Environments: Challenges and the Quest for Inhibition. Antibiotics (Basel) 10(2021).

10. Ho, C.M. et al. High rate of qacA- and qacB-positive methicillin-resistant Staphylococcus aureus isolates from chlorhexidine-impregnated catheter-related bloodstream infections. Antimicrob Agents Chemother 56, 5693–7 (2012).

11. Reddy, V.S., Shlykov, M.A., Castillo, R., Sun, E.I. & Saier, M.H., Jr. The major facilitator superfamily (MFS) revisited. FEBS J 279, 2022–35 (2012).

12. Drew, D., North, R.A., Nagarathinam, K. & Tanabe, M. Structures and General Transport Mechanisms by the Major Facilitator Superfamily (MFS). Chem Rev 121, 5289–5335 (2021).

13. Brawley, D.N. et al. Structural basis for inhibition of the drug efflux pump NorA from Staphylococcus aureus. Nat Chem Biol (2022).

14. Heng, J. et al. Substrate-bound structure of the E. coli multidrug resistance transporter MdfA. Cell Res 25, 1060–73 (2015).

15. Li, X.Z., Zhang, L. & Nikaido, H. Efflux pump-mediated intrinsic drug resistance in Mycobacterium smegmatis. Antimicrob Agents Chemother 48, 2415–23 (2004).

16. Brown, M.H. & Skurray, R.A. Staphylococcal multidrug efflux protein QacA. J Mol Microbiol Biotechnol 3, 163–70 (2001).

17. Tennent, J.M. et al. Physical and biochemical characterization of the qacA gene encoding antiseptic and disinfectant resistance in Staphylococcus aureus. J Gen Microbiol 135, 1–10 (1989).

18. Grkovic, S., Brown, M.H., Roberts, N.J., Paulsen, I.T. & Skurray, R.A. QacR is a repressor protein that regulates expression of the Staphylococcus aureus multidrug efflux pump QacA. J Biol Chem 273, 18665–73 (1998).

19. Schumacher, M.A. et al. Structural basis for cooperative DNA binding by two dimers of the multidrug-binding protein QacR. EMBO J 21, 1210–8 (2002).

20. Mitchell, B.A., Paulsen, I.T., Brown, M.H. & Skurray, R.A. Bioenergetics of the staphylococcal multidrug export protein QacA. Identification of distinct binding sites for monovalent and divalent cations. J Biol Chem 274, 3541–8 (1999).

21. Majumder, P. et al. Dissection of Protonation Sites for Antibacterial Recognition and Transport in QacA, a Multi-Drug Efflux Transporter. J Mol Biol 431, 2163–2179 (2019).

22. Kumar, S. et al. Structural basis of inhibition of a transporter from Staphylococcus aureus, NorC, through a single-domain camelid antibody. Commun Biol 4, 836 (2021).

23. Zomot, E. et al. A New Critical Conformational Determinant of Multidrug Efflux by an MFS Transporter. J Mol Biol 430, 1368–1385 (2018).

24. Kumar, S., Mahendran, I., Athreya, A., Ranjan, R. & Penmatsa, A. Isolation and structural characterization of a Zn(2+)-bound single-domain antibody against NorC, a putative multidrug efflux transporter in bacteria. J Biol Chem 295, 55–68 (2020).

25. Martinez Molledo, M., Quistgaard, E.M., Flayhan, A., Pieprzyk, J. & Low, C. Multispecific Substrate Recognition in a Proton-Dependent Oligopeptide Transporter. Structure 26, 467–476 e4 (2018).

26. Debruycker, V. et al. An embedded lipid in the multidrug transporter LmrP suggests a mechanism for polyspecificity. Nat Struct Mol Biol (2020).

27. Nagarathinam, K. et al. Outward open conformation of a Major Facilitator Superfamily multidrug/H(+) antiporter provides insights into switching mechanism. Nat Commun 9, 4005 (2018).

28. Wu, J., Hassan, K.A., Skurray, R.A. & Brown, M.H. Functional analyses reveal an important role for tyrosine residues in the staphylococcal multidrug efflux protein QacA. BMC Microbiol 8, 147 (2008).

29. Zhang, X.C., Zhao, Y., Heng, J. & Jiang, D. Energy coupling mechanisms of MFS transporters. Protein Sci 24, 1560–79 (2015).

30. Del Alamo, D., Sala, D., McHaourab, H.S. & Meiler, J. Sampling alternative conformational states of transporters and receptors with AlphaFold2. Elife 11(2022).

31. Del Alamo, D., Govaerts, C. & McHaourab, H.S. AlphaFold2 predicts the inward-facing conformation of the multidrug transporter LmrP. Proteins 89, 1226–1228 (2021).

32. Smirnova, I. et al. Transient conformers of LacY are trapped by nanobodies. Proc Natl Acad Sci U S A 112, 13839–44 (2015).

33. Abramson, J. et al. Structure and mechanism of the lactose permease of Escherichia coli. Science 301, 610–5 (2003).

34. Smirnova, I. et al. Outward-facing conformers of LacY stabilized by nanobodies. Proc Natl Acad Sci U S A 111, 18548–53 (2014).

35. Hassan, K.A. et al. Optimized production and analysis of the staphylococcal multidrug efflux protein QacA. Protein Expr Purif 64, 118–24 (2009).

36. Ahmed, S., Bhasin, M., Manjunath, K. & Varadarajan, R. Prediction of Residue-specific Contributions to Binding and Thermal Stability Using Yeast Surface Display. Front Mol Biosci 8, 800819 (2021).

37. Chattopadhyay, G. et al. Functional and Biochemical Characterization of the MazEF6 Toxin-Antitoxin System of Mycobacterium tuberculosis. J Bacteriol 204, e0005822 (2022).

38. Punjani, A., Rubinstein, J.L., Fleet, D.J. & Brubaker, M.A. cryoSPARC: algorithms for rapid unsupervised cryo-EM structure determination. Nat Methods 14, 290–296 (2017).

39. Punjani, A., Zhang, H. & Fleet, D.J. Non-uniform refinement: adaptive regularization improves single-particle cryo-EM reconstruction. Nat Methods 17, 1214–1221 (2020).

40. Terwilliger, T.C., Ludtke, S.J., Read, R.J., Adams, P.D. & Afonine, P.V. Improvement of cryo-EM maps by density modification. Nat Methods 17, 923–927 (2020).

41. Pettersen, E.F. et al. UCSF Chimera--a visualization system for exploratory research and analysis. J Comput Chem 25, 1605–12 (2004).

42. Emsley, P. & Cowtan, K. Coot: model-building tools for molecular graphics. Acta Crystallogr D Biol Crystallogr 60, 2126–32 (2004).

43. Afonine, P.V. et al. Real-space refinement in PHENIX for cryo-EM and crystallography. Acta Crystallogr D Struct Biol 74, 531–544 (2018).

44. Wu, E.L. et al. CHARMM-GUI Membrane Builder toward realistic biological membrane simulations. J Comput Chem 35, 1997–2004 (2014).

45. Jo, S., Kim, T., Iyer, V.G. & Im, W. CHARMM-GUI: a web-based graphical user interface for CHARMM. J Comput Chem 29, 1859–65 (2008).

46. Lee, J. et al. CHARMM-GUI Input Generator for NAMD, GROMACS, AMBER, OpenMM, and CHARMM/OpenMM Simulations Using the CHARMM36 Additive Force Field. J Chem Theory Comput 12, 405–13 (2016).

