## Supplementary figure 1-6 for "CryoEM structure of QacA, an antibacterial efflux transporter from *Staphylococcus aureus*"

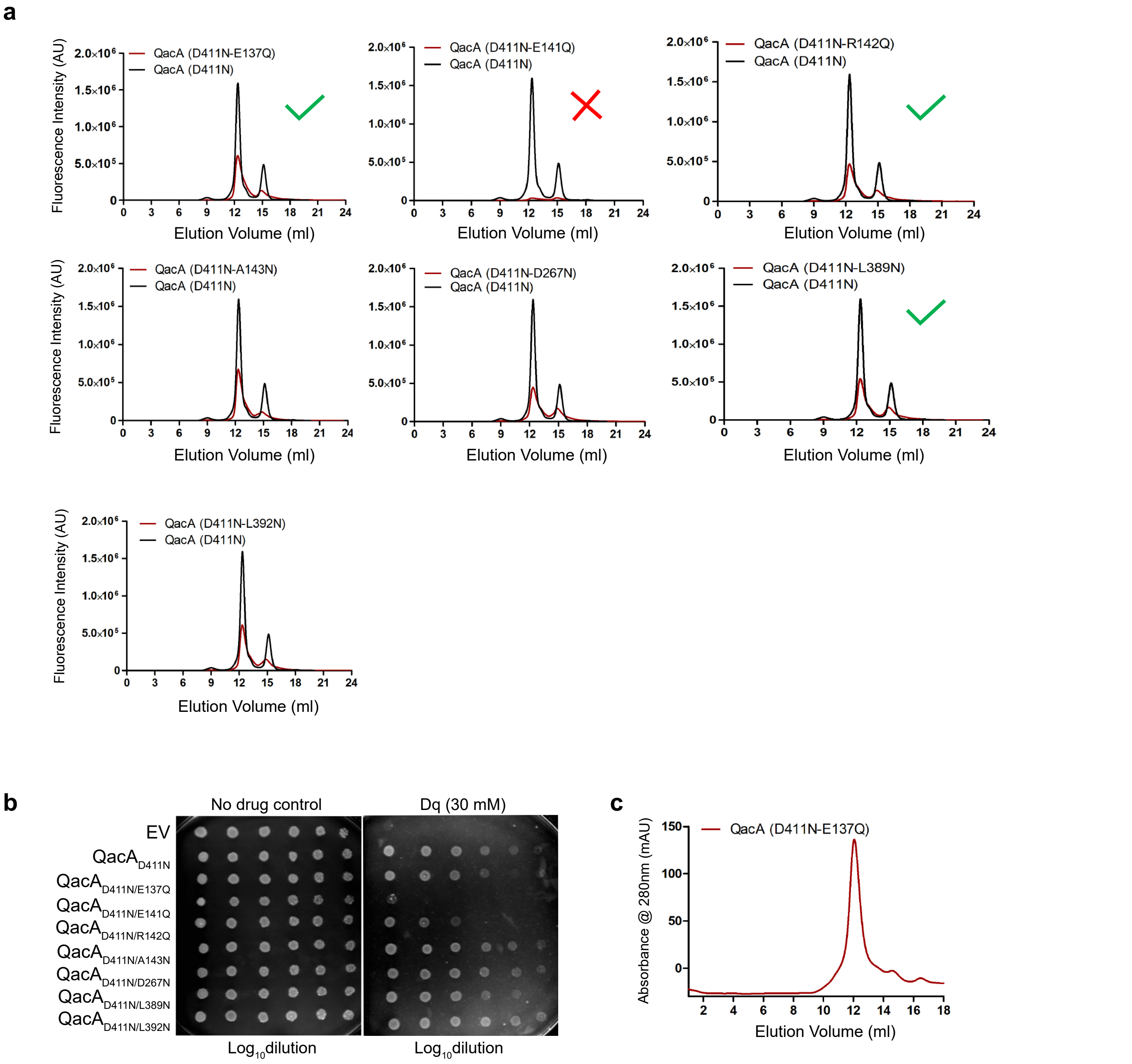


**Supplementary Fig. 1 | Screening of QacA mutants for immunization. a,** FSEC profiles of all the cytosolic rim residue mutants screened and **b,** Survival assay on JD838 cells to screen for stable functional construct of QacA. Constructs with monodisperse FSEC profile but with reduction in function (i.e., E137Q, E142Q and L389N in the background of D411N) were shortlisted as such constructs could be more stable in one conformation and out of these, D411N-E137Q was chosen randomly to proceed with as antigen for immunization. EV = empty vector. Mutations of QacA WT are labelled against the serial dilutions in **b**. **c,** SEC profile of $QacA_{D411N/E137Q}$ mutant.

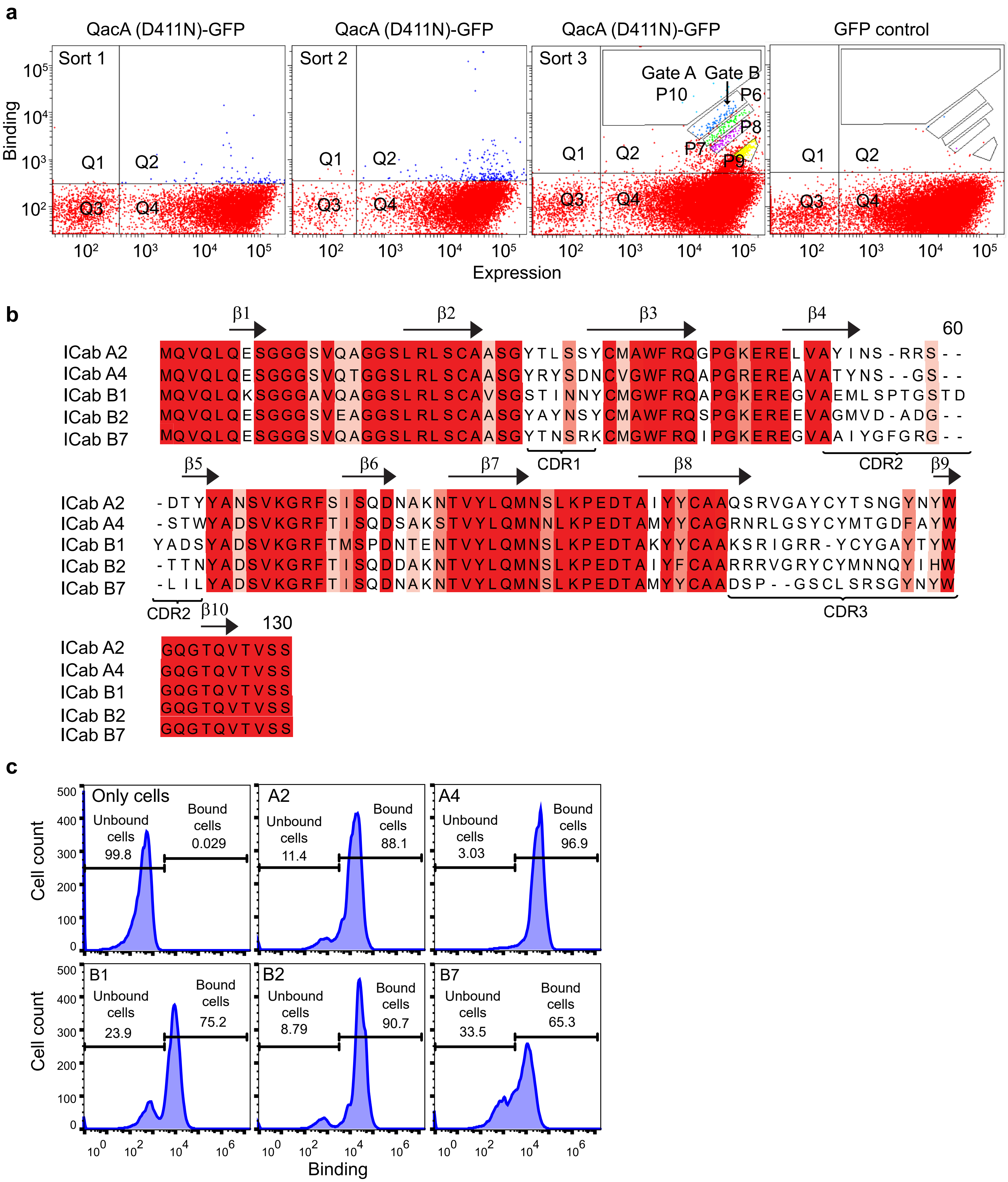


**Supplementary Fig. 2 | Five unique ICabs isolated against QacA$_{D411N}$. a,** FACS enrichment of ICab binders against QacA through sequential sorting of Q2 from Sort 1 to Sort 3. ICabs were isolated from Gates A (P10) and B (P6). **b,** Multiple sequence alignment of ICabs A2 and A4 (from P10), and B1, B2 and B7 (from P6) with higher sequence conservations highlighted with saturated shades of red. **c,** Population shifts of cells expressing ICabs A2, A4, B1, B2 or B7 on their surface in the presence of micellar QacA$_{D411N\text{-}GFP}$. Bound and unbound fractions are represented in percentages.

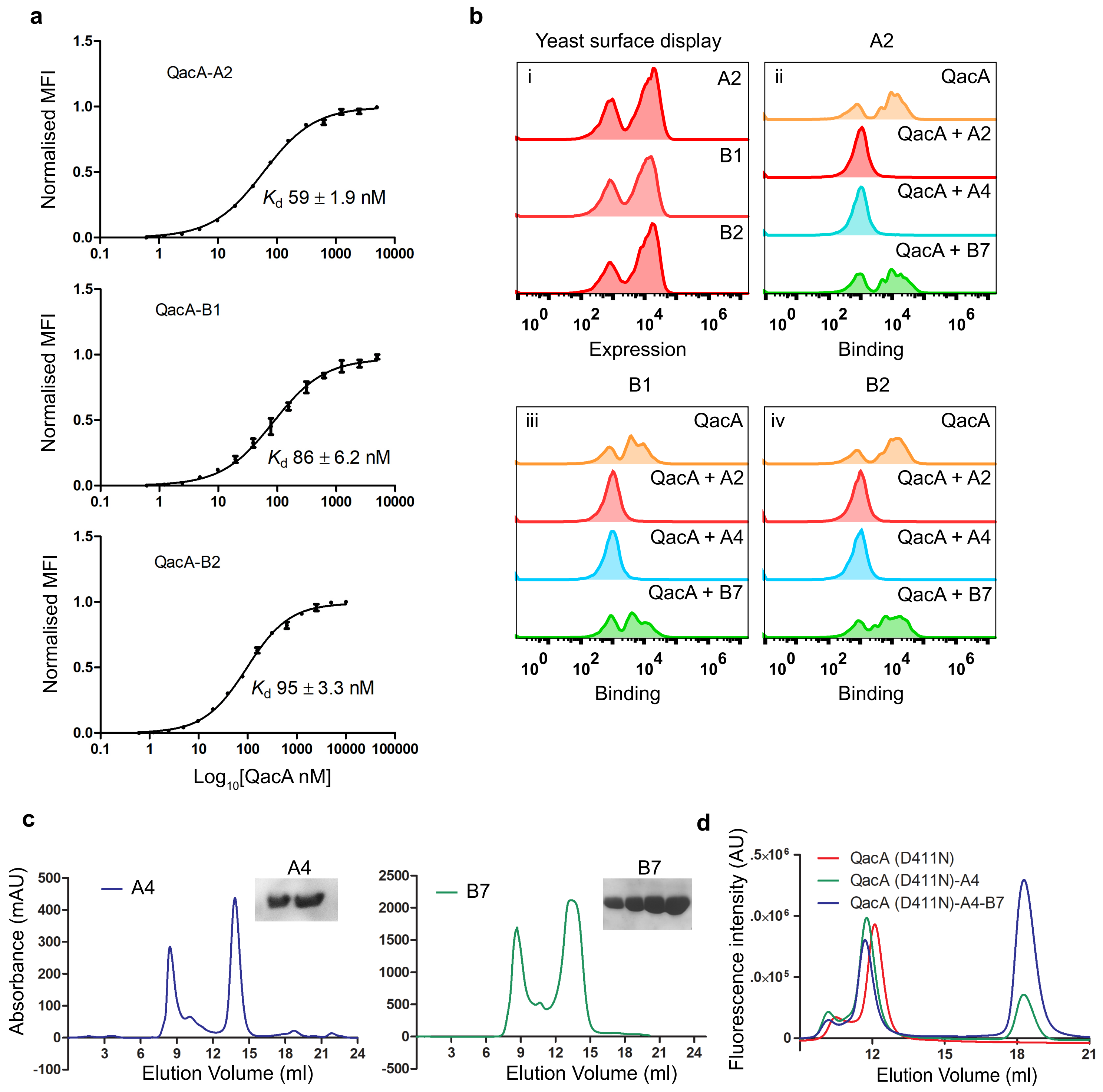


**Supplementary Fig. 3 | Characterization of affinities of ICabs for QacA. a,** Flow cytometry derived affinity measurements of ICabs A2 (top), B1 (middle) and B2 (bottom) for QacA. Error bars represent SEM with n=3 for technical triplicates. **b,** Population shift based titrations of ICabs against QacA-GFP preincubated with other ICabs in the group. A rightward shift of population infers non-competitive binding of ICabs. **c,** SEC profiles of ICabs A4 (left) and B7 (right). Collected fractions around 13-14 ml of elution volume were run on SDS-PAGE and the corresponding bands are shown (inset). **d,** FSEC profiles with leftward shifts in elution volumes of $QacA_{D411N}$ in the presence of A4 and B7 indicate increase in apparent molecular mass of QacA upon binding with either or both the ICabs.

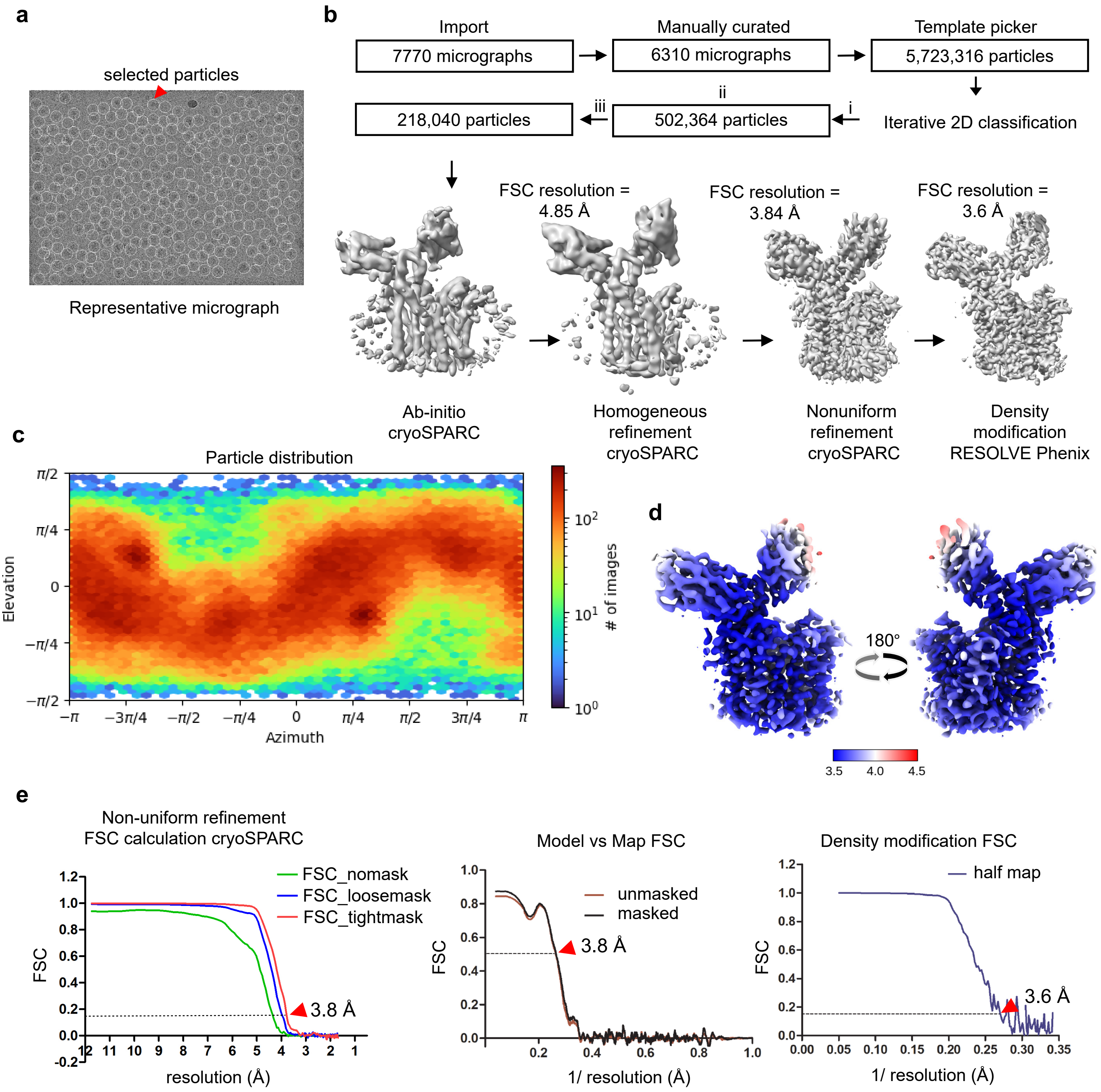


**Supplementary Fig. 4 | Workflow for structural determination of QacA-ICabs complex through cryoEM.** **a,** Representative micrograph from the 7770 movies collected. White rings highlight the particles picked for further processing. **b,** Summarised workflow of the data processing in CryoSPARC. Each process written is followed below by a value for micrographs/particles used in that step (boxed). For iterative 2D class averaging, 5,723,316 particles were picked in iteration (i), followed by 502,364 and 218,040 particles in iteration (ii) and (iii) respectively. **c,** 2D heat map of particle orientation distribution in the processed data. **d,** Local resolution maps of QacA-ICabs complex in the final structure. **e,** Fourier shell correlation plots of the final dataset with resolutions pointed at FSCs of 0.143 and 0.5 for non-uniform refinement and density modification, and model versus map FSC comparison respectively.

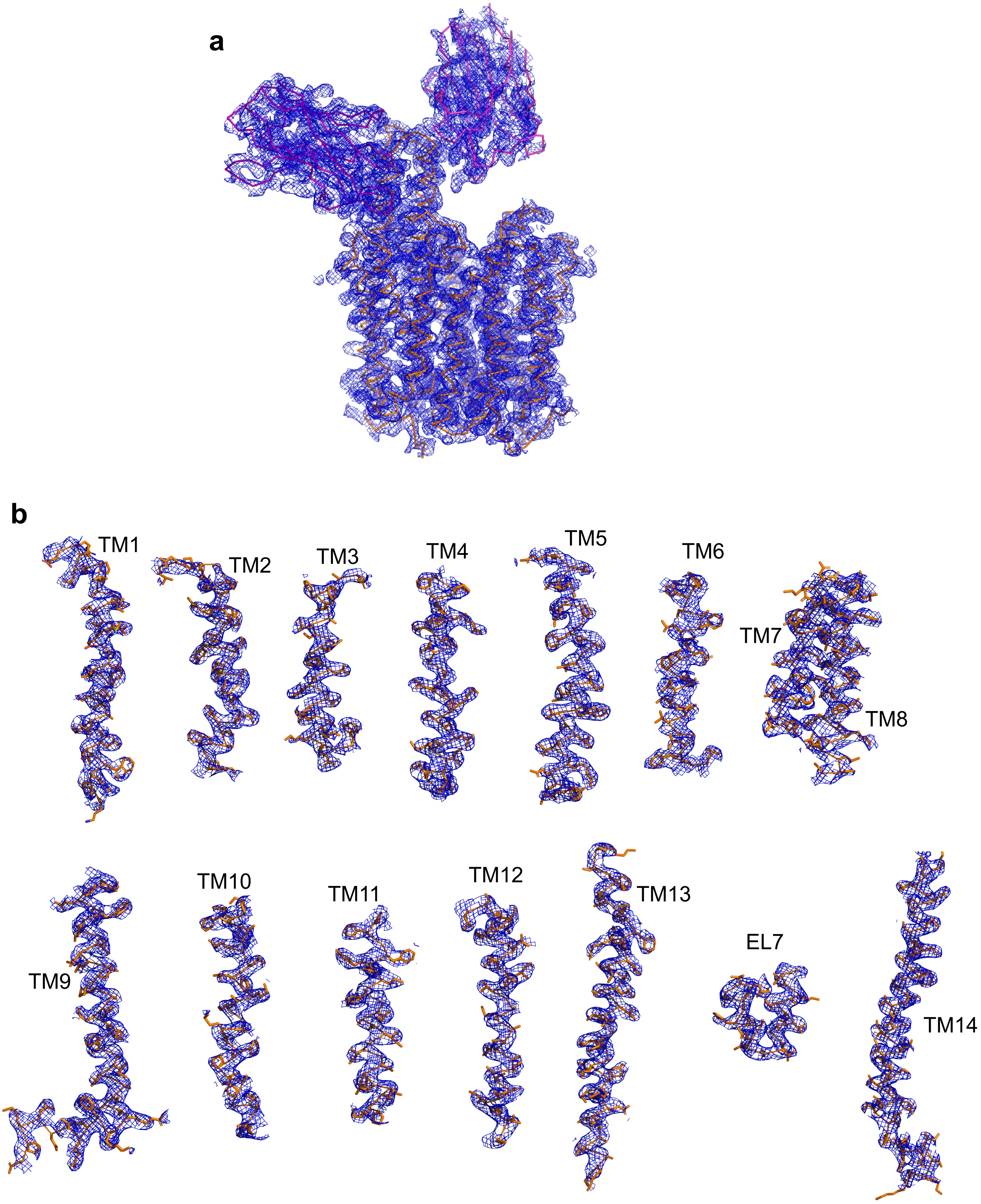


**Supplementary Fig. 5 | Model fit for QacA-ICab complex and individual TM helices. a,** Coulomb potential map for QacA-ICabs complex contoured at 2σ with **b,** individual transmembrane helices modelled.

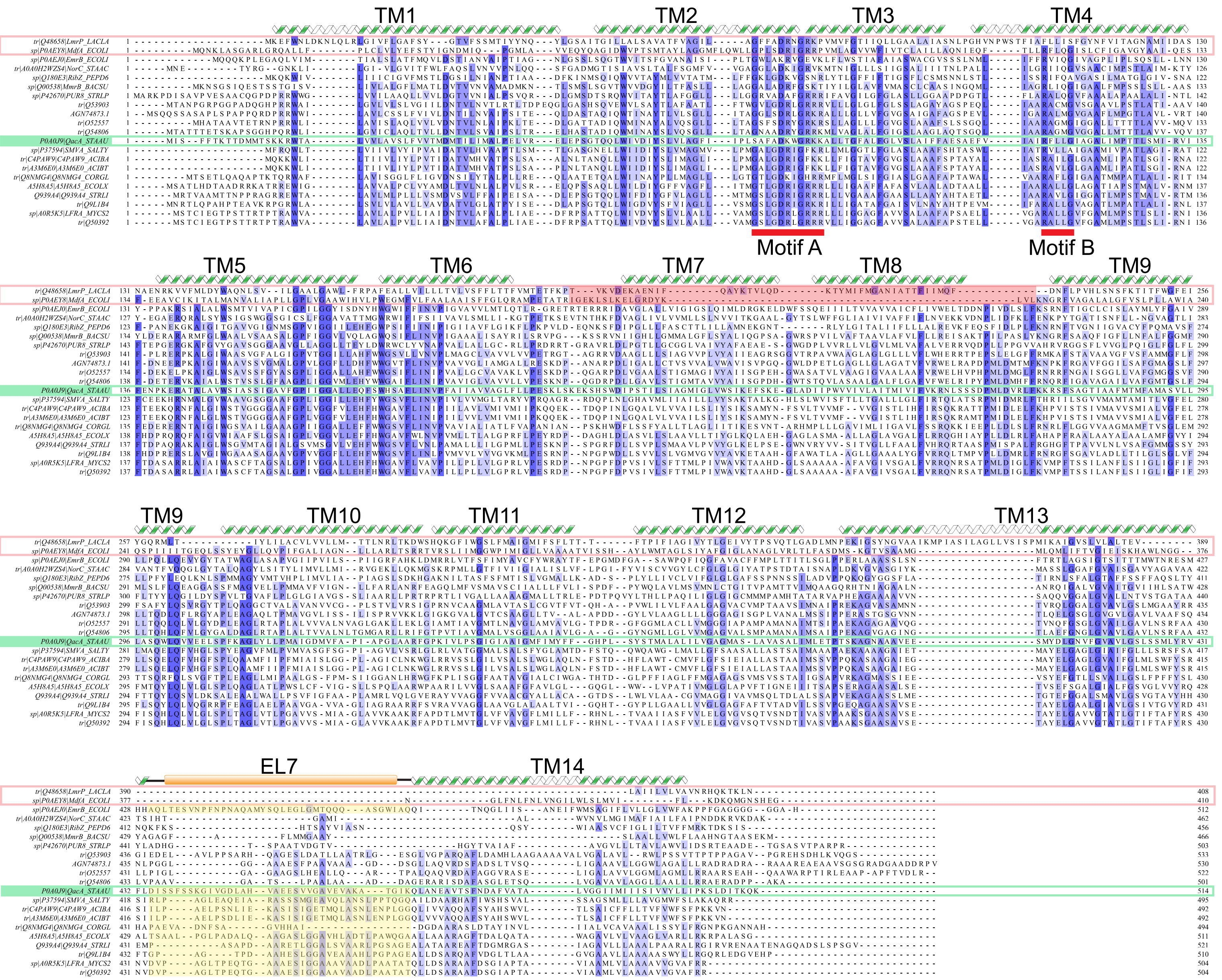


**Supplementary Fig. 6 | Sequence and structural alignments of QacA with DHA1/2 transporters.** Multiple sequence alignment of QacA with related sequences from different prokaryotic genera. QacA is highlighted in green and DHA1 transporters MdfA and LmrP are boxed in red. Extents for transmembrane helices of QacA are shown above the MSA. Sequences for horizontal linker helix between TMs 6 and 7 in MdfA and LmrP, that substitute for linker TMs 7 and 8 in DHA2 transporters are highlighted in red, while the extracellular loop 7 (EL 7) sequences in QacA homologs are highlighted in yellow. The conserved Motif A is demarcated with a red underline.

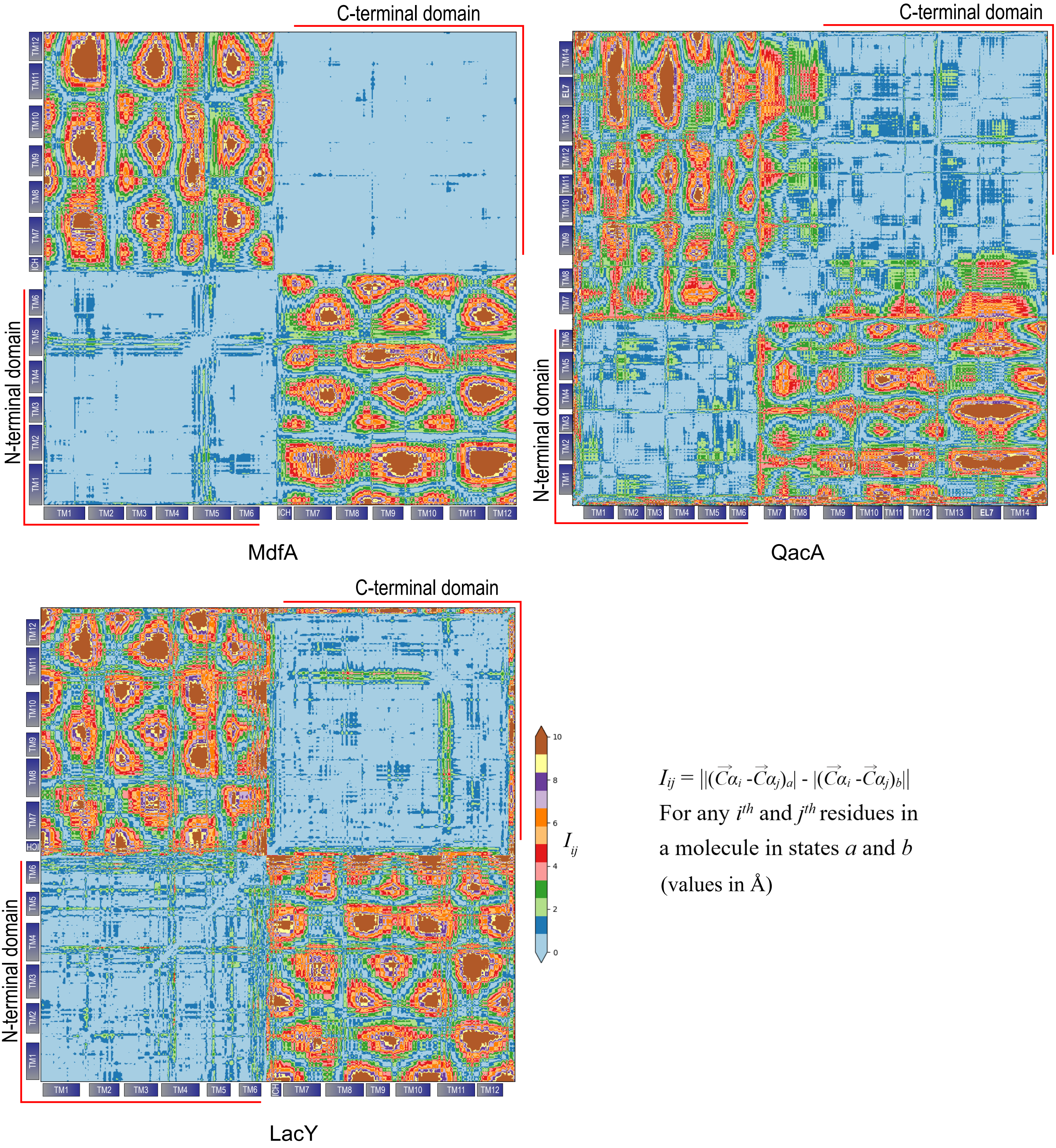


**Supplementary Fig. 7 | Asymmetric rocker switch mechanism of transport in QacA.** Difference plot of Cα vs C α distances of transporters in outward open and inward open states. Values are in Angstroms. Colour scheme is annotated besides the plot of LacY, with emphasis on the differences lying between 0 to 10 Å; distances greater than 10 Å have been annotated with the same colour for the sake of clarity. For QacA, $\text{QacA}_{\text{D411N}}$ is taken as the outward open state and $\text{QacA}_{\text{io}}$ model has been taken for the inward open state. Similarly, PDBIDs 4ZP0 and 6GV1 were used for inward and outward open states respectively of MdfA, and PDBIDs 1PV7 and 5GXB were used for inward and outward open states respectively for LacY. Residues which are modelled in both the states of a transporter are analysed. Extents of individual TMs and N- and C-terminal domains are annotated.
