## Supplementary data table 1 for "CryoEM structure of QacA, an antibacterial efflux transporter from *Staphylococcus aureus*"

Supplementary Table 1. CryoEM data collection, refinement and validation.

| Parameters | QacA-ICab complex EMDB (EMD-33612) PDB id (7Y58) |
|---|---|
| **Data Collection/Processing** | |
| Microscope | FEI Titan Krios |
| Voltage (kV) | 300 |
| Detector | K3 Quantum |
| Magnification | 105,000 |
| Defocus range | -1.0 – -4.0 (μm) |
| Pixel size | 0.831 |
| Electron exposure ($e^-$/Å$^2$) | 51 |
| Exposure time | 2 |
| Symmetry Imposed | C1 |
| Initial particle number | 5,723,316 |
| Final particle number | 218,040 |
| Map Resolution | 3.6 |
| FSC threshold | 0.143 |
| Map resolution | |
| **Refinement** | |
| Initial model | QacA Alphafold2 model |
| Map resolution (masked) (FSC 0.143) | 3.8 Å |
| Map resolution (Density modification) (FSC 0.143) | 3.6 Å |
| **Model Composition** | |
| Non-Hydrogen atoms | 5609 |
| Protein residues | 743 |
| Ligands | - |
| **RMS deviations** | |
| Bond lengths | 0.003 |
| Bond Angles | 0.748 |
| **Validation** | |
| Refined model CC | 0.77 |
| Molprobity score | 2.12 |
| Clashscore | 11.3 |
| **Ramachandran plot** | |
| Favoured | 92 |
| Allowed | 8 |
| Disallowed | 0 |
